# Raver1 links *Ripk1* RNA splicing to caspase-8-mediated pyroptotic cell death, inflammation, and pathogen resistance

**DOI:** 10.1101/2024.11.27.625707

**Authors:** Boyao Zhang, Pontus Orning, Jesse W. Lehman, Alexandre Dinis, Leslie Torres-Ulloa, Roland Elling, Michelle A. Kelliher, John Bertin, Megan K. Proulx, Liv Ryan, Richard Kandasamy, Terje Espevik, Athma A. Pai, Katherine A. Fitzgerald, Egil Lien

## Abstract

Multiple cell death and inflammatory signaling pathways converge on two critical factors: receptor interacting serine/threonine kinase 1 (RIPK1) and caspase-8. Careful regulation of these molecules is critical to control apoptosis, pyroptosis and inflammation. Here we discovered a pivotal role of Raver1 as an essential regulator of *Ripk1* pre-mRNA splicing, expression, and functionality, and the subsequent caspase-8-dependent inflammatory cell death. Macrophages from *Raver1*-deficient mice exhibit altered splicing of *Ripk1*, accompanied by diminished cell death and reduced activation of caspase-8, Gasdermin D and E, caspase-1, as well as decreased interleukin-18 (IL-18) and IL-1ß production. These effects were triggered by *Yersinia* bacteria, or by restraining TAK1 or IKKβ in the presence of LPS, TNF family members, or IFNγ. Consequently, animals lacking *Raver1* showed heightened susceptibility to *Yersinia* infection. Raver1 and RIPK1 also controlled the expression and function of the C-type lectin receptor Mincle. Our study underscores the critical regulatory role of Raver1 in modulating innate immune responses and highlights its significance in directing *in vivo* and *in vitro* inflammatory processes.

**Significance:** Caspase-8 and the kinase RIPK1 are at focal points of several inflammation and cell death pathways. Thus, a careful regulation of their actions is needed. Our work identifies the RNA splicing factor Raver1 as a critical factor directing the splicing of *Ripk1* in order to modulate RIPK1/caspase-8-driven pyroptosis, apoptosis and inflammation. Raver1 is central for macrophage responses to *Yersinia* bacteria, initiated after blockade of kinases TAK1 and IKK, measured as activation of RIPK1, caspase-8, Gasdermin D, caspase-3, IL-1ß and IL-18. Importantly, Raver1 is necessary for host resistance to *Yersinia* infection *in vivo*. We propose that Raver1 is key for correct tuning of RIPK1-caspase-8 dependent processes.

## Introduction

Cell death and inflammatory pathways frequently share signaling mechanisms critical for inflammation during microbial infection or tissue injury. Receptor interacting serine/threonine kinase 1 (RIPK1) is a key regulator in both inflammatory pathways and apoptosis, pyroptosis, and necroptosis pathways. RIPK1 is engaged upstream of caspase-8 activation during extrinsic apoptotic cascades and is required for optimal nuclear factor-kappa-B (NF-κB) signaling after lipopolysaccharide (LPS) or tumor necrosis factor (TNF) stimulation (1, 2). Therefore, RIPK1 and caspase-8 are integral mediators governing innate immune signaling processes in inflammation and antimicrobial host defenses. RIPK1 is tightly regulated via phosphorylation, ubiquitylation, and proteolytic cleavage (3). Additional mechanisms likely influence RIPK1-caspase-8-dependent pathways. Further investigations into these intricate regulatory mechanisms are crucial to understanding the pathophysiology of many inflammatory conditions to develop targeted therapeutic interventions.

Essential information on the interconnectivity between cell death and inflammatory pathways has come from studies using *Yersinia* bacteria as a model system. Human-pathogenic *Yersinia* bacteria include *Y. pestis*, the causative agent of plague, and the gastrointestinal pathogens *Y. pseudotuberculosis* and *Y. enterocolitica*. During *Yersinia* infection, the bacterial Type III secretion system effector YopJ (or YopP in *Y. enterocolitica*) inhibits TGF-ß-activated kinase 1 (TAK1), the IκB kinase complex (IKKα/ß), and mitogen-activated protein kinases through acetylation of phosphorylation sites (4–7). Both TAK1 (8, 9) and IKKα/β (10, 11) are essential to restrain RIPK1-driven cytotoxicity and inflammatory pathologies via inhibitory phosphorylation of RIPK1. Thus, *Yersinia* infection unleashes RIPK1 kinase activity and the subsequent assembly of cytosolic cell death complex II, involving RIPK1, FADD, and caspase-8 (1, 2, 12, 13). This pathway licenses apoptosis by activating caspase-8 (13, 14). We and others have recently presented evidence for the crosstalk between apoptotic and pyroptotic pathways. In immune cells, *Yersinia*-activated caspase-8 also cleaves Gasdermin D (GSDMD), initiating atypical pyroptosis and allowing the release of pro-inflammatory cytokines interleukin-1ß (IL-1ß) and interleukin-18 (IL-18) (2, 15–17), as well as inducing secondary caspase-1 cleavage (13–15), which further amplifies inflammation.

*Yersinia*-triggered RIPK1-caspase-8-dependent cell death is characterized by a complex interplay between components of apoptosis and pyroptosis pathways. The effect of YopJ can also be mimicked by treatment with LPS or TNF combined with pharmacological inhibition of TAK1 (TAK1-i) or IKK (IKK-i), whereas the impact of interferon-γ (IFNγ) in the context of TAK1 blockade is unclear. YopJ-induced cell death is independent of caspase-1, whereas the caspase-8-dependent release of IL-1ß is partially influenced by caspase-8-controlled caspase-1 activation (15). Loss of *Gsdmd* delays cell death upon blockade of TAK1 or IKKα/ß and reduces IL-1ß and IL-18 secretion (15, 17).

Using a CRISPR-based screen combined with *Yersinia* infection, we identified Raver1 as a novel regulator of RIPK1-caspase-8 signaling by influencing pre-mRNA splicing of *Ripk1*.

Loss of *Raver1* promotes the inclusion of an alternative *Ripk1* exon, resulting in the production of a truncated dysfunctional neo-peptide and reduced overall RIPK1 levels. *Raver1*-deficient macrophages and neutrophils display impaired responses to *Yersinia* or LPS/TNF/IFNγ+TAK1-blockade-induced pyroptosis, and IL-1ß/IL-18 release. Importantly, compromised anti-microbial responses culminated in the decreased ability of *Raver1*-deficient animals to resist *Yersinia* challenge. We also extend these findings and establish that Raver1 regulates C-type lectin receptor (CLR) Mincle (*Clec4e*) expression and signaling via its effects on RIPK1. Together, our study defines a critical role for Raver1 in regulating innate immune responses, emphasizing its importance in orchestrating inflammatory processes both *in vitro* and *in vivo*.

## Results

### A CRISPR screen in primary macrophages identifies Raver1 as a regulator of *Yersinia*-induced RIPK1-caspase-8-mediated cell death

To identify new regulators of RIPK1-caspase-8-mediated signaling, we performed an unbiased genome-wide CRISPR-Cas9 cell death screen using primary mouse bone marrow-derived macrophages (BMDM) infected with *Yersinia*. In order to better focus on the caspase-8 signaling component, bone marrow cells from Cas9-expressing *Rip3^−/−^* mice were transduced with a genome-wide single-guide RNA (sgRNA) library and differentiated into macrophages. Once progenitor cells were fully differentiated, cells were infected with *Y. pestis*. DNA from the surviving populations was isolated, and enriched sgRNAs were identified. The screen identified several known components, including RIPK1, GSDMD, and caspase-8, as well as components of the lysosomal Rag-Ragulator complex (Figure 1A), recently described as important regulators of RIPK1-caspase-8 activation (18). In addition, top hits included two proteins involved in alternative splicing: Raver1 and Ptbp1 (Figure 1A). Raver1 and Ptbp1 are both implicated in pre-mRNA splicing regulation, including alternative splicing of α-tropomyosin exon 3 in smooth muscle cells (19–21). Raver1 is proposed to cooperate with Ptbp1, the latter exerts a broad influence on a wide range of splicing events (22), and its deletion is associated with embryonic mortality in mice. The specific contribution of Raver1 to innate immune function *in vivo* and *in vitro* however remains largely unexplored. Thus, we focused additional analysis on the newly identified hit—Raver1.

**Figure 1.**
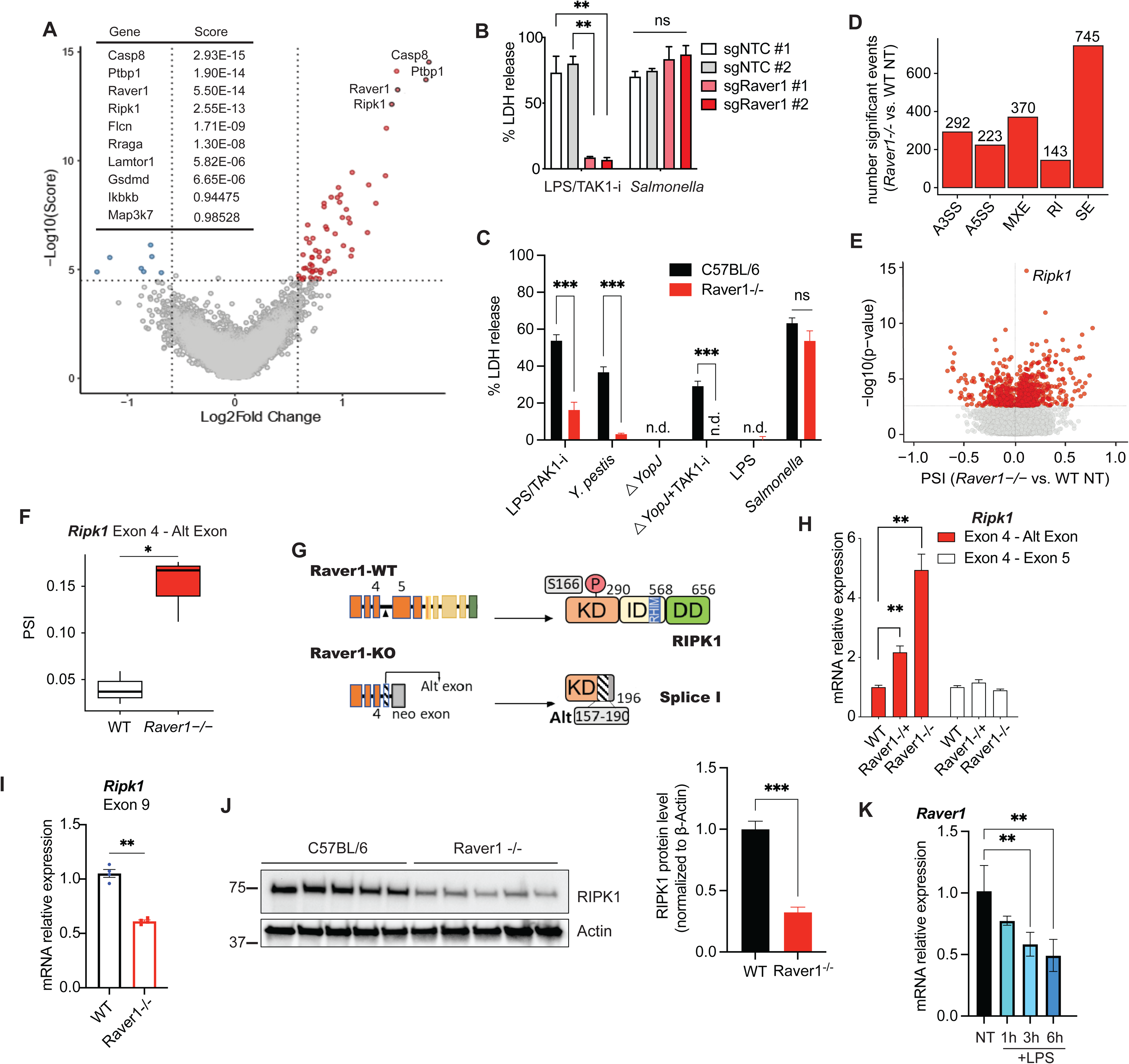
Raver1 regulates RIPK1 RNA splicing and signaling. (**A**) A CRISPR screen was performed in Cas9-expressing *Ripk3*^−/−^ bone marrow-derived macrophages (BMDMs) infected with *Yersinia pestis*. Log_2_-fold change and MAGeCK RRA enrichment scores are shown to indicate the essentiality of the genes. (**B, C**) Cas9-expressing BMDMs transduced with *Raver1*-targeting sgRNAs or a non-targeting control (NTC) (B) or *Raver1^−/−^* BMDMs (C) were challenged with *Y. pestis*, *Y. pestis*△YopJ (△YopJ) or LPS and TAK1 inhibitor (TAK1-i), or *Salmonella*. Cell death was measured by lactate dehydrogenase (LDH) release after 4 hours. (**D**) Frequencies of significant alternative splicing events in BMDMs from *Raver1^−/−^ vs.* WT mice, as reported by rMATS differential splicing analysis conducted using RNA-seq data in three independent BMDM replicates per genotype. A3SS, alternative 3′ splice sites; A5SS, alternative 5′ splice sites; MXE, mutually exclusive exons; RI, retained intron; SE, skipped exons (**E**) Volcano plot depicting skipped exon alternative splicing events detected by rMATS analysis, with percent spliced-in (PSI) of *Raver1^−/−^* SE events *vs.* WT SE events on the x-axis, and – log_10_(p-value) reported by rMATS on the y-axis. *Ripk1* is highlighted as the most significant skipped exon change upon *Raver1* deletion. The horizontal dotted line corresponds to the false discovery rate (FDR) significance threshold of 10%, with points ≤ 10% FDR threshold colored red. **(F**) *Ripk1* alternative exon 4 inclusion in WT vs. *Raver1^−/−^* BMDMs, with percent spliced-in (PSI) for alternative exon 4 detected by rMATS on the y-axis. (**G**) Proposed model for canonical and alternative splicing of *Ripk1* in the presence or absence of *Raver1.* Briefly, inclusion of an alternative exon between exon 4 and 5 resulted in production of a subsequent protein variant Splice I, preserving the majority of kinase domain (KD), but lacking auto-phosphorylation site S166, intermediate domain (ID), RIP homotypic interaction motif (RHIM), and death domain (DD). (**H, I**) mRNA levels of *Ripk1* exons in indicated resting BMDMs as quantified by qPCR. (**J**) Protein levels of RIPK1 in WT or *Raver1^−/−^* BMDMs from 5 mice; the right-hand side shows quantification from the Western blot. (**K**) mRNA levels of *Raver1* in C57BL/6 BMDMs treated with LPS for the indicated times were quantified by qRT-PCR. Data are presented as the mean ± SD (B, C, K) or ± SEM (H-J) for triplicates from three or more independent experiments. (H-K) data normalized to β-Actin. For comparison between two groups, Student’s t-test was used; for more than two groups, ANOVA with Bonferroni post-hoc test was used. ns, not significant; *p≤0.05; **p≤0.01; ***p≤0.001.*Abbreviations*: n.d., not detected; NT, non-treated; Alt exon, alternative exon.

We first confirmed the effect of Raver1 by transducing Cas9 transgenic primary BMDMs with sgRNAs targeting *Raver1*. Compared to control cells transduced with non-targeting sgRNAs, cells lacking *Raver1* were protected from cell death induced by *Yersinia* infection or by mimicking *Yersinia* infection using the combined action of LPS plus TAK1-i (Figures 1B and S1A). This inhibition was specific to RIPK1-caspase-8-induced death as cells lacking *Raver1* responded normally to NLRC4-caspase-1-mediated pyroptosis induced in response to *Salmonella enterica* serovar Typhimurium (Figure 1B) (23).

To determine the role of Raver1 in innate immunity, we generated mice lacking *Raver1* via CRISPR/Cas9. This yielded a line (#6) with a 171 bp deletion in exon 1 of the *Raver1* gene which led to a loss of any detectable protein expression (Figures S1B-S1D). Unlike *Ptbp1^−/−^* mice that are embryonically lethal, *Raver1^−/−^*mice are viable and healthy throughout life expectancy.

BMDMs from these mice recapitulated protection from *Yersinia*-or chemical-mimetic-induced cytotoxicity found in sg*Raver1*-transduced cells (Figure 1C).

### Raver1 represses alternative splicing of *Ripk1* to maintain RIPK1 protein abundance

Given the proposed role of Raver1 in the regulation of alternative splicing, we sought to determine how the loss of *Raver1* reshapes the transcriptome and contributes to *Yersinia-*induced cytotoxicity. We performed differential splicing analyses on RNA-seq datasets in untreated BMDMs from *Raver1^−/−^ vs.* WT mice, revealing pervasive global changes in transcriptome composition (Figures 1D and S1D). Of the 1773 significant alternative splicing events, 745 were skipped exon events (Figure 1D) and there was a strong bias toward skipped exon inclusion in *Raver1^−/−^* BMDMs (Figures 1D and S1E). Together, these data indicate that Raver1 is important for repressing exon inclusion during pre-mRNA splicing.

Strikingly, a *Ripk1* alternative exon shows the most significant skipped exon change upon *Raver1* deletion (Figure 1E). Indeed, *Ripk1* was the only key member of the caspase-8 signaling pathway found to have significant Raver1-mediated splicing events. Thus, these analyses identified *Ripk1* as a potential splicing target of Raver1 in primary macrophages, whose dysregulation in the absence of *Raver1* might modulate *Yersinia*-induced cytotoxicity.

Interestingly, Ptbp1 was proposed as a regulator of necroptosis in a fibroblast cell line by impacting *Ripk1* alternative splicing (24), and additional cell death screen also suggested involvement of Raver1 (25). Therefore, we hypothesized that Raver1 functions in caspase-8-mediated innate immune responses by modulating the splicing of *Ripk1* in primary cells.

An in-depth investigation on *Ripk1* splicing patterns in BMDMs validated our findings that loss of *Raver1* altered *Ripk1* transcript composition, through increased inclusion of the *Ripk1* alternative exon between exon 4 and exon 5 (Figures 1F and S1F), causing a frameshift that introduced an early stop codon. We predicted this would result in a truncated RIPK1 isoform variant with a protein neo-sequence at the C-terminus, that we called Splice I (Figure 1G). Although Splice I shares sequences with the majority of the RIPK1 kinase domain, it lacks a key portion of it — phosphorylation site Ser166, as well as crucial domains, including intermediate, RIP homotypic interaction motif (RHIM), and death domains.

We designed qPCR assays quantifying *Ripk1*canonical exons 4-5, full-length *Ripk1* (exemplified by exon 9), and the inclusion of the *Ripk1*alternative exon (exon 4-alt. exon), linked to the production of Splice I. This analysis suggested that *Raver1^−/−^*BMDMs had a five-fold increase in Splice I expression, as measured by the inclusion of the alternative exon, compared to WT BMDMs, as well as an almost 50% reduction in full-length *Ripk1* transcripts as a consequence (Figures 1H, 1I and S1G). Accordingly, full-length RIPK1 protein was reduced by over 60% in the absence of *Raver1* (Figures 1J and S1H). This reduction can potentially be attributed to nonsense-mediated decay of transcripts harboring the premature stop codon. Of note, the RIPK1 antibody recognizes the C-terminus of the protein and cannot detect the splice variant. In contrast to RIPK1, *Raver1*^−/−^ macrophages or *Raver1* sgRNA-treated cells had comparable protein levels of caspase-8, TAK1, FADD, and RIPK3 (Figures S1H and S1I).

Importantly, expression of *Raver1* appeared to decrease with time following stimulation with LPS in macrophages (Figure 1K). This indicates the presence of a feedback loop that could limit excessive inflammation in order to achieve homeostasis following pathway activation.

### Raver1 is required for RIPK1-caspase-8-mediated cell death in primary phagocytes

To examine how Raver1 mediates cell death, we infected WT and *Raver1*^−/−^ cells with *Yersinia*. BMDMs from *Raver1*^−/−^ mice were significantly protected from pyroptosis induced by human-pathogenic *Yersinia* spp. *(Y. pestis, Y. pseudotuberculosis, Y. enterocolitica)* (Figure 2A), while maintaining a normal response to *Salmonella* (Figure 2B). Additionally, BMDMs from *Raver1^−/−^*mice were protected from pyroptosis following treatments that mimic *Yersinia* infection, including TAK1 or IKK inhibitors in combination with LPS, TNF, or its family member lymphotoxin-α (LT-α) (Figures 2B-2D, S2A, and S2B). Macrophages from a second *Raver1*-deficient mouse line (#4) (Figures S1B and S1C) had a similar phenotype (Figure S2C). The delayed death in *Raver1*^−/−^ BMDMs was also seen in peritoneal macrophages (Figure S2D). Recent studies have revealed that in neutrophils, RIPK1 regulates pyroptosis and IL-1 release through Gasdermin E (GSDME) during *Yersinia* infection (26). Interestingly, we observed attenuated cell death in *Raver1*^−/−^ neutrophils following TNF+TAK1-i treatment (Figure 2E), indicating a broader role of Raver1 in different immune cells involved in innate immunity.

**Figure 2.**
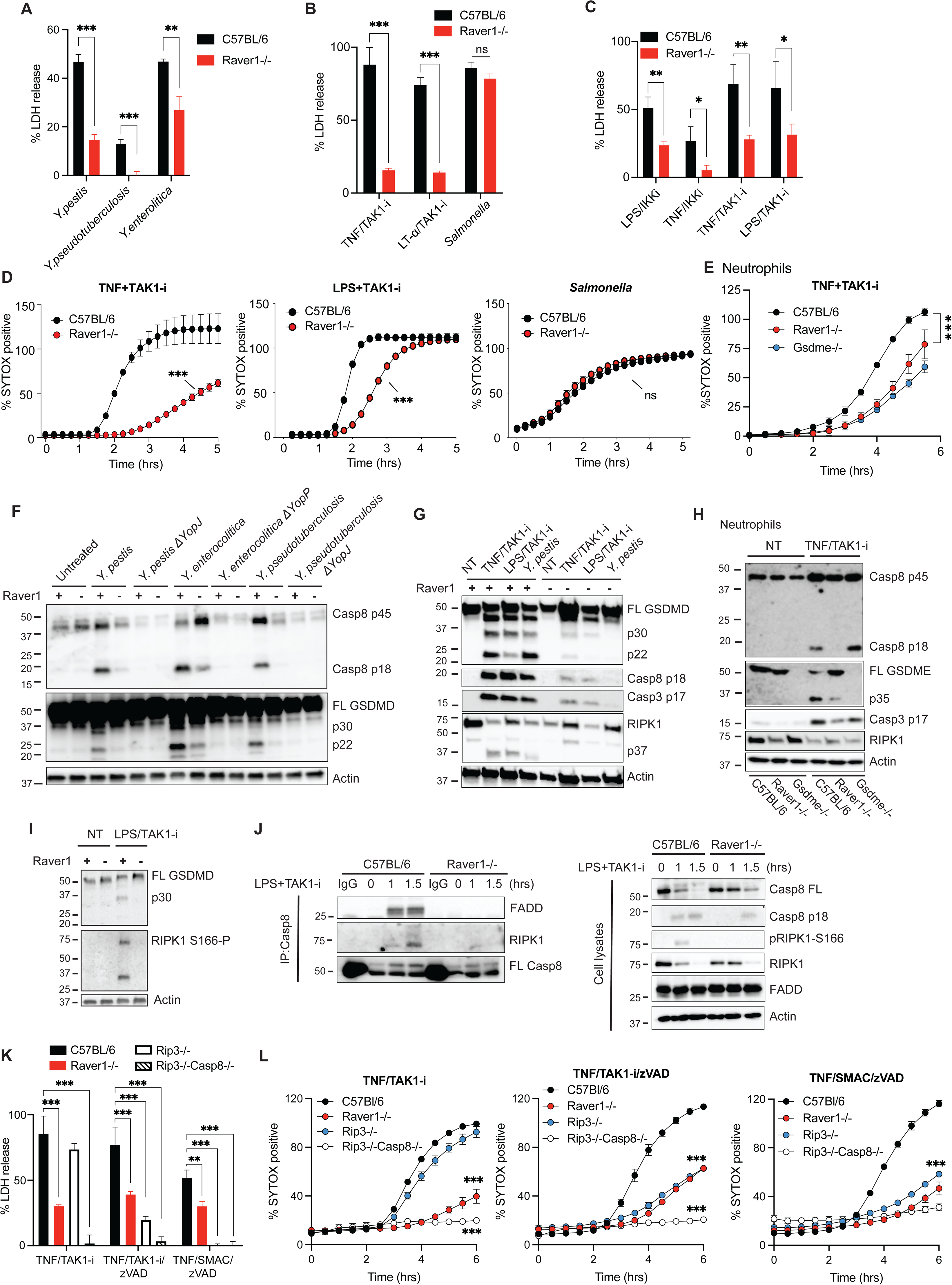
*Raver1*-deficient macrophages and neutrophils are resistant to pyroptotic cell death and activation of RIPK1, caspase-8, and GSDMD/E after *Yersinia* challenge. (**A–E**) C57BL/6 or *Raver1*^−/−^ BMDMs (A–D) or neutrophils (E) from indicated mice were challenged with *Yersinia* spp., *Salmonella*, TAK1-i or IKK inhibitor (IKK-i) plus LPS, TNF, or lymphotoxin-α (LT-α). Cell death was measured by LDH release after 4 hours (A–C) or SYTOX dye membrane permeability (D, E). (F–I) Cell lysates from WT, *Raver1*^−/−^ BMDMs (F, G, I) or neutrophils (H) from indicated mice were analyzed by immunoblotting for GSDMD/E, caspase-8, caspase-3, and RIPK1 after 2–3 hours (F–H) or 1 hour (I). (**J**) Caspase-8 immunoprecipitation (IP: Casp8) or cell lysates of C57BL/6 or *Raver1*^−/−^ BMDMs treated with LPS+TAK1-i for 0, 1 or 1.5 hours were analyzed by immunoblotting on FADD, RIPK1 and caspase-8. (**K, L**) Indicated BMDMs were stimulated with TNF, TAK1-i, or a SMAC mimetic, in the presence or absence of a pan-caspase inhibitor zVAD-fmk pretreatment, then analyzed for LDH release after 4 hours (K) or SYTOX dye membrane access (L). Data are presented as the mean ± SD for triplicate wells from three or more independent experiments. Immunoblots are representative of ≥3 performed. For comparison between two groups, Student’s t-test was used; for more than two groups, ANOVA with Bonferroni post-hoc test was used. n.s., not significant; *p≤0.05, **p≤0.01, ***p≤0.001. *Abbreviations*: FL, full-length; NT, non-treated; +, *Raver1*^+/+^; –, *Raver1*^−/−^ (F, G, I).

Deleting MLKL did not further protect against cell death induced by TNF or LPS plus TAK1-i in *Raver1*-deficient macrophages (Figure S2E), suggesting that no secondary activation of the necroptosis pathway was occurring following this pyroptotic/apoptotic response.

In macrophages, caspase-8 cleaves GSDMD into a pore-forming p30 N-terminal fragment to trigger pyroptosis, and subsequent caspase-3 activation contributes to additional processing to a p22 fragment (15). *Raver1*^−/−^ BMDMs displayed reduced caspase-8 and GSDMD processing in response to *Yersinia* spp. in a YopJ/YopP-dependent manner (Figure 2F).

Furthermore, activation of GSDMD and caspase-8 was also compromised in *Raver1*^−/−^ macrophages when the infection was mimicked by TNF, LT-α, or LPS plus TAK1-i treatment (Figures 2G, S2F, and S2G). Thus, the protection from cell death observed in *Raver1*^−/−^ BMDMs was due to reduced processing and activation of caspase-8 and GSDMD.

We next examined other proteins downstream of caspase-8 and observed blunted processing of RIPK1, caspase-3, and GSDME in Raver1-deficient macrophages and neutrophils (Figures 2G, 2H, and S2H), events all correlating with diminished RIPK1-caspase-8-mediated cell death. This aligns with a recent study proposing that RIPK1 regulates GSDMD and GSDME processing in macrophages and neutrophils (26). Moreover, *Raver1*^−/−^ cells did not show any detectable phosphorylation of RIPK1 S166, after TAK1-blockade with LPS or TNF family members (Figures 2I and S2G), suggesting RIPK1 activation is greatly suppressed in the absence of *Raver1*. This aligns with the proposed model of Splice I lacking the RIPK1 S166 activation site, a key autophosphorylation residue (27). In *Raver1*^−/−^ cells, limited RIPK1-caspase-8 association was detected following LPS+TAK1 inhibition (Figure 2J), indicating that formation of cell death complex II is defective in *Raver1*-deficient BMDMs. This likely reflects a reduction of the critical complex component, RIPK1. Together, this information points toward a model in which Raver1 regulates RIPK1 upstream of RIPK1-caspase-8 death complex assembly, likely by balancing the levels of full-length RIPK1 and the Splice I variant.

RIPK1 kinase is also required for controlling RIPK3-MLKL-dependent necroptosis upon caspase inhibition (28). Similar to *Rip3^−/−^* BMDMs, deletion of *Raver1* significantly decreased susceptibility to necroptosis induced by pan-caspase inhibitor zVAD-FMK in addition to TNF and TAK1-i or SMAC mimetic (Figures 2K and 2L). This is consistent with a previous observation that CRISPR-RNA-mediated knock-out of Ptbp1 downregulates necroptosis in fibroblast cells (24). Taken together, the above evidence demonstrates that Raver1 is indispensable for RIPK1-caspase-8-mediated pyroptosis/apoptosis as well as RIPK1-RIPK3-mediated necroptosis.

### Raver1-deficient primary phagocytes display impaired inflammatory responses

Our prior work indicated that this RIPK1-caspase-8-GSDMD pathway is directly involved in *Yersinia*-induced release of IL-1ß and IL-18, and these pro-inflammatory cytokines are important for host resistance to *Yersinia* strains (13, 15, 29–31). Reduced RIPK1-caspase-8-dependent release of IL-1ß was observed in *Raver1*^−/−^ macrophages and *Raver1*^−/−^ neutrophils following *Yersinia* infection, or TNF/LPS plus TAK1 or IKK inhibition (Figures 3A-3D, S3A-C). This reduction in IL-1ß correlated with reduced activation of caspase-8 and GSDMD/E in *Raver1*^−/−^ cells (Figures 2F-2I and S2F-S2H).

**Figure 3.**
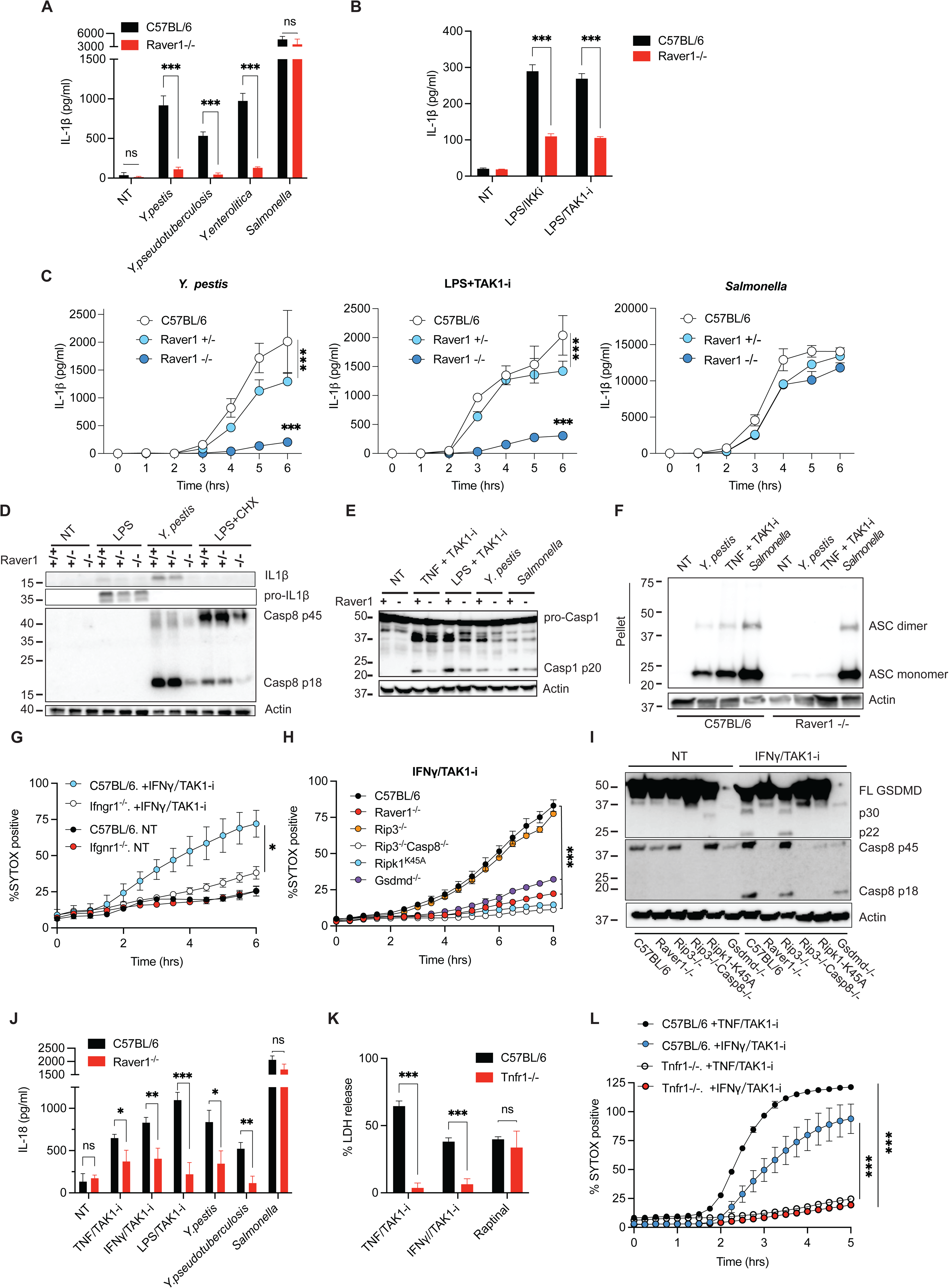
Raver1 controls TAK1-restrained IL-1β/IL-18 release, and IFNγ-driven inflammatory responses. (**A-C**) C57BL/6 or *Raver1*^−/−^ BMDMs were treated with *Yersinia* spp., *Salmonella*, or LPS with TAK1-i or IKK-i. IL-1β release into the supernatant as measured by ELISA after 5 hours or at the indicated time points. (**D–E**) Cell lysates plus supernatants were analyzed by immunoblotting for caspase-1, –8, or IL-1β cleavage 3 hours after the indicated stimulation. (**F**) Oligomerization of ASC in the inflammasome-enriched and crosslinked lysates as detected by immunoblotting. (**G–I**) BMDMs of indicated genotypes were treated with IFNγ plus TAK1-i. Cell death was measured by SYTOX dye membrane permeability (G, H). GSDMD and caspase-8 processing at 3 hours were detected by immunoblotting (I). (**J**) IL-18 levels as measured by ELISA on supernatants from C57BL/6 or *Raver1*^−/−^ BMDMs challenged with indicated bacteria or TAK1-i for 5 hours. (**K, L**) C57BL/6 or *Tnfr1*^−/−^ BMDMs were treated with TAK1-i plus TNF or IFNγ, or Raptinal alone. Cell death was measured by LDH release after 4.5 hours (K), or SYTOX dye membrane permeability (L). Data are presented as the mean ± SD for triplicates from three or more independent experiments. Immunoblots are representative of 3 immunoblots performed. For comparison between two groups, Student’s t-test was used; for more than two groups, ANOVA with Bonferroni post-hoc test was used. n.s., not significant; *p≤0.05, **p≤0.01, ***p≤0.001. *Abbreviations*: CHX, cycloheximide.

Inhibition and deletion of TAK1 (8, 15) or IKKβ (32, 33) is associated with IL-1β release and ASC-NLRP3 inflammasome activity. We have previously suggested that ASC-NLRP3 functions downstream of caspase-8 and potassium efflux in these stimulatory conditions (15).

Consequently, caspase-1 p20 cleavage and ASC inflammasome oligomerization were also abrogated in *Raver1*^−/−^ macrophages (Figures 3E, 3F, and S3D). Furthermore, our data suggest that the absence of *Raver1* in BMDMs does not directly influence IL-1ß secretion induced by other inflammasome ligands that trigger activation of caspase-1-dependent inflammasomes: NLRP3 (Pam3CSK4+ATP or nigericin), NLRC4 (*Salmonella*), Pyrin (Pam3CSK4+TcdB toxin and *Y. pestis* ΔYopMΔYopJ), and AIM2 (Pam3CSK4 + poly[dA:dT]) (Figures S3E and S3F). Taken together, Raver1 controls RIPK1-caspase-8-dependent release of IL-1ß and secondary ASC inflammasome activation, contributing to inflammation.

Another key cytokine for immune responses and antimicrobial defenses is interferon-γ (IFNγ). IFNγ levels are elevated during bacterial and viral infections, including *Yersinia* and SARS-CoV-2 (29, 34). Less is known about cytotoxicity involving IFNγ, but recent evidence has implicated IFNγ in inducing RIPK1-caspase-8-mediated cell death and inflammation during the cytokine storm and multi-organ failure following SARS-CoV-2 challenge (34). Furthermore, IFNγ+LPS stimulation also engages caspase-8 (35), whereas the role of RIPK1 in this condition is unclear. Strikingly, our experiments showed that IFNγ+TAK1-i also induced considerable cell death in WT BMDMs (Figures 3G and S4A, S4B). In contrast, *Raver1*^−/−^ macrophages were protected against IFNγ+TAK1-i-induced cytotoxicity (Figures 3H and S4C, S4D) with a concomitant reduction in caspase-8, GSDMD, GSDME, and caspase-3 activation (Figures 3I and S4E, S4F). The cell death was controlled by RIPK1, GSDMD, and caspase-8 (Figures 3H and S4C, S4G). IFNγ+TAK1-i stimulation appeared fundamentally different from IFNγ+LPS in that the latter condition did not involve RIPK1 kinase activity or Raver1 (Figure S4G).

IL-18 is an IFNγ-inducing factor, and both cytokines play critical roles in host defenses during *Yersinia* infection (29, 31). We found that IFNγ+TAK1-i induces the release of a substantial amount of IL-18 protein in a Raver1-dependent manner, similar to *Yersinia* infection, or TAK1-i plus LPS or TNF (Figure 3J). In contrast, Raver1 did not influence IL-18 induced by *Salmonella* (Figure 3J), consistent with the lack of impact of Raver1 on *Salmonella*-triggered IL-1ß and cell death (Figures 2 and 3).

To determine how inflammation impacts IFNγ-induced cell death, we examined TNFR1 signaling. We previously reported a partial role for TNFR1 in *Yersinia*-or LPS+TAK1-i-induced macrophage death (15), we now observed that IFNγ+TAK1-i-induced cytotoxicity was heavily influenced by TNF signaling, as the absence of TNFR1 reduced cell death to background levels (Figures 3K and 3L). These data suggest that TAK1 activity restrains IFNγ-induced cell death and inflammation via secondary TNF release and TNFR1 signaling, mediated by Raver1-regulated RIPK1 and caspase-8.

### The Splice I variant interacts with RIPK1 to negatively regulate cell death

To further explore the role of TNF and confirm the role of Raver1 and RIPK1 splice variants in a different system, we used Cas9-expressing WEHI164-clone13 fibrosarcoma cells, a highly TNF-sensitive cell line that has been used in a death-based bioassay for TNF activity (36). We observed that these cells underwent RIPK1 and caspase-mediated cell death following treatment with TNF or TNF+TAK1-i, abolished by deletion of *Raver1* (Figures 4A and S5A).

**Figure 4.**
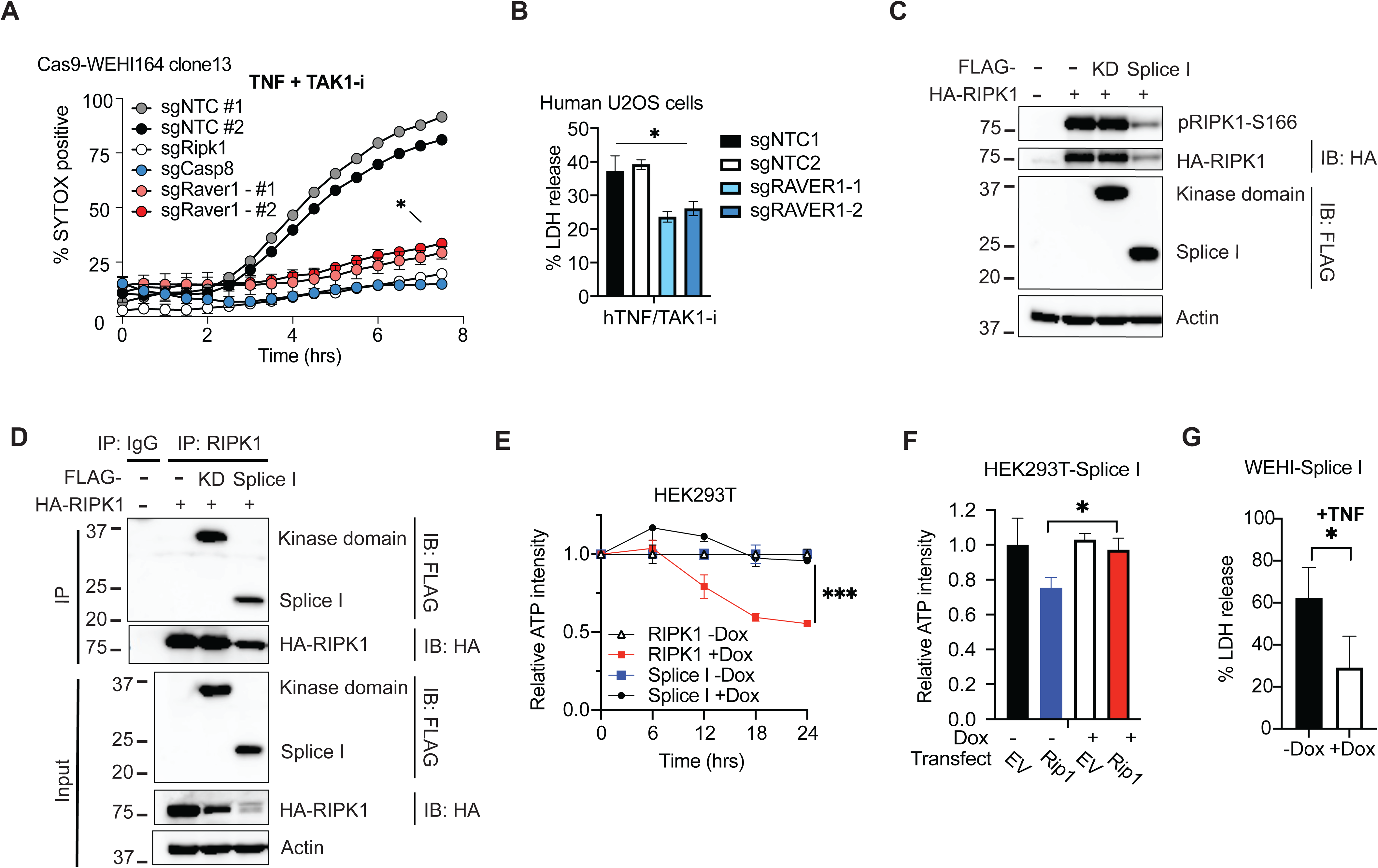
Raver1 dysfunction results in a RIPK1 splicing isoform, Splice I, that negatively regulates cell death. (**A, B**) WEHI164 clone 13 fibrosarcoma cells (A) or human U2OS osteosarcoma cells (B) transduced with indicated sgRNAs were treated with TNF and TAK1-i. Cell death was measured by SYTOX dye membrane permeability (A) or LDH release after 10 hours (B). (**C, D**) HEK293T cells were co-transfected with vectors encoding RIPK1-HA and RIPK1 variants (Splice I-Flag or RIPK1-kinase domain [KD]-Flag) or empty pcDNA3 for 24 hours. Cell lysates were analyzed by immunoblotting (C), or immunoprecipitated with RIPK1 antibodies followed by immunoblotting against Flag or HA (D). (**E**) Doxycycline-induced RIPK1 or RIPK1-Splice I was expressed in HEK293T cells. Cell death is measured by reduced ATP intensity after doxycycline (Dox) was added for the indicated times. (**F**) Doxycycline-induced RIPK1-Splice I was expressed in HEK293T cells, with doxycycline addition 12 hours before transfection of RIPK1 or empty vector (EV). Cell death is measured by reduced ATP intensity after 12 hours. (**G**) Doxycycline-induced RIPK1-Splice I was expressed in WEHI164 clone 13 cells, with doxycycline addition 12 hours before stimulation with TNF (50 ng/ml). Cell death was measured by LDH release after 10 hours. Data are presented as the mean ± SD for triplicates from three or more independent experiments. Immunoblots are representative of 3 immunoblots performed. For comparison between two groups, Student’s t-test was used; for more than two groups, ANOVA with Bonferroni post-hoc test was used. ns, not significant; *p≤0.05, ***p≤0.001. *Abbreviations*: NTC, non-targeting control; hTNF, human TNF;.

These effects were conserved in human cells, as sgRNAs against *Raver1* also reduced TNF+TAK1-i-induced cell death mediated by RIPK1/Caspase-8 in human U2OS osteosarcoma cells (Figures 4B, S5B and S5C).

To study the function of the RIPK1 Splice I variant (Figure 1G), we ectopically expressed FLAG-tagged Splice I in HEK293T cells, and this resulted in the production of a 22 kDa peptide (Figure 4C), indicating Splice I can be translated into protein. Splice I interacts with full-length RIPK1, as evidenced by the detection of HA-tagged RIPK1 from immunoprecipitates of FLAG-Splice I when co-expressed (Figure 4D). This was accompanied by reduced expression and activation of newly synthesized RIPK1 (HA-tagged) (Figures 4C and 4D). However, we did not observe a reduction in pre-existing RIPK1 protein utilizing a doxycycline-induced Splice I system (Figure S5D). This suggests that increased expression of Splice I has a negative effect on the new synthesis of RIPK1 protein. The reduced RIPK1 levels could be a key aspect in the downregulation of signaling.

The Splice I variant was unable to drive cytotoxicity alone but reduced RIPK1-induced cytotoxicity (Figures 4E-4F). Data in Fig 5A indicated that WEHI164 cells could be suitable for testing Splice I function. Ectopically expression of the RIPK1 Splice I product in WEHI164-clone13 cells reduced TNF-induced cell death (Figure 4G). These results could be explained by a dominant-negative action of Splice I binding to full-length RIPK1, and consequently, reduced ability of RIPK1 to induce cell death. Taken together, these findings suggest that Raver1 regulates RIPK1-caspase-8-mediated cell death and inflammation through alternative splicing of *Ripk1*. Loss of *Raver1* reduced full-length RIPK1 levels and increased expression of a dysfunctional truncated Splice I variant that bound to RIPK1 and inhibited its ability to induce cell death. Collectively, these events have profound effects on normal RIPK1 function.

**Figure 5.**
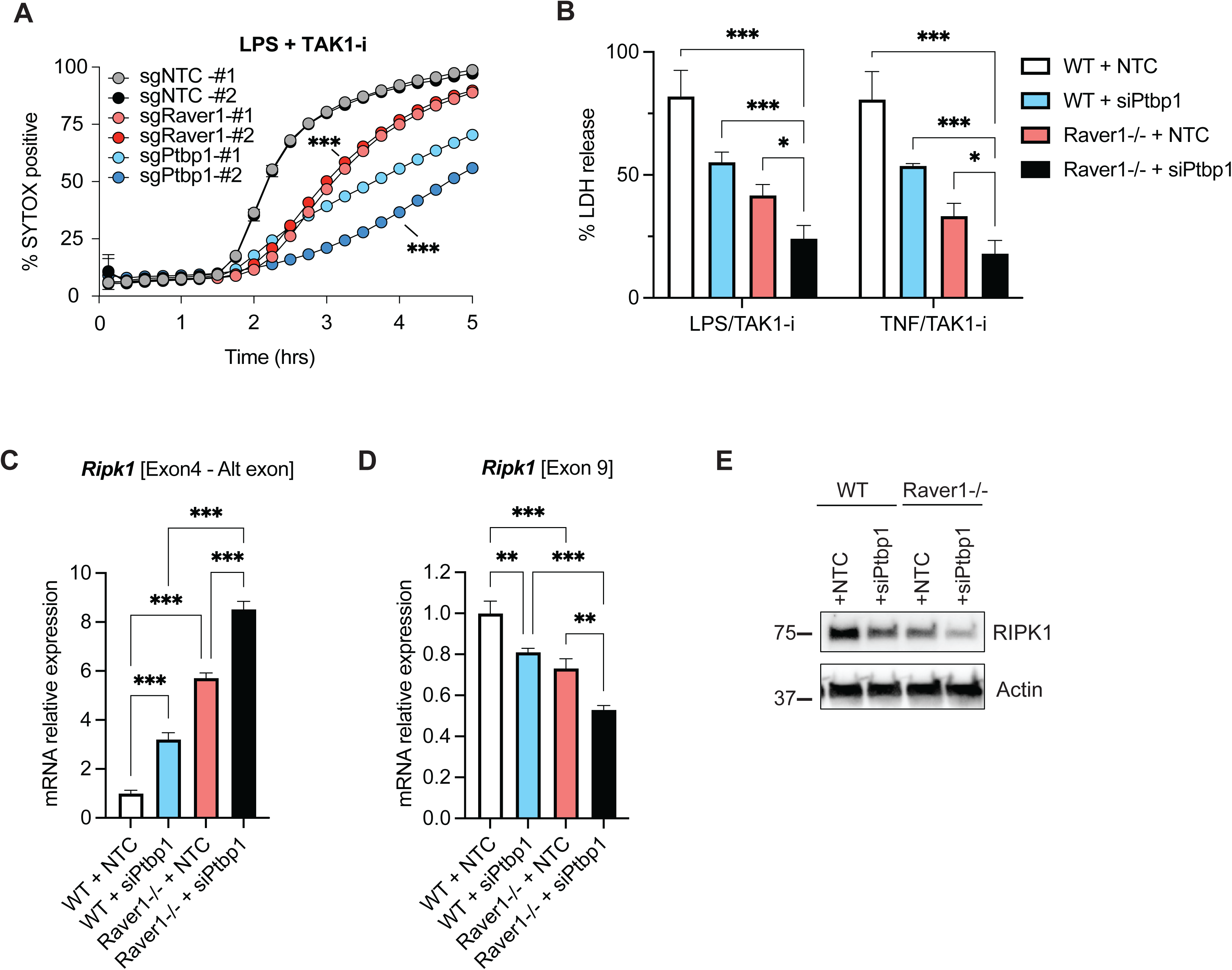
Raver1 cooperates with Ptbp1 to regulate RIPK1 splicing and cell death. (**A**) Cas9-expressing BMDMs transduced with sgRNA targeting *Raver1* or *Ptbp1* were treated with LPS plus TAK1-i and cell death was measured by SYTOX dye membrane permeability. (**B-E**) WT or *Raver1^−/−^*immortalized macrophages were transfected with *Ptbp1*-targeting siRNA or non-targeting control (NTC) siRNA. Cell death was measured by LDH release 4 hours after LPS+TAK1-i or TNF+TAK1-i treatment (B). mRNA levels of *Ripk1* exons were quantified by qRT-PCR, normalized to *Actin* (C, D). Protein levels of RIPK1 were analyzed by immunoblotting (E). Data are presented as the mean ± SD for triplicate wells from three or more independent experiments. Immunoblots are representative of three blots performed. For comparison, ANOVA was used with Bonferroni’s post-hoc test. *p≤0.05, **p≤0.01, ***p≤0.001. *Abbreviations*: Alt exon, alternative exon.

### Raver1 cooperates with Ptbp1 to regulate *Ripk1* splicing and cell death

Raver1 has been proposed as a Ptbp1 co-factor in regulating alternative splicing events but may also function independently of Ptbp1 (20, 37). Both Raver1 and Ptbp1 ranked high on our CRISPR screen (Figure 1A), and deletion of either by CRISPR/Cas9 was associated with increased inclusion of the *Ripk1* alternative exon and diminished cytotoxicity in Cas9-BMDMs treated with LPS/TAK1-i (Figures 5A, S1A, S6A and S6B). These data suggested that both Raver1 and Ptbp1 could influence exon exclusion in *Ripk1*. To test the cooperative effects of the two splicing factors, we transfected WT or *Raver1^−/−^*macrophages with siRNA against *Ptbp1* (Figure S6C). *Ptbp1* knockdown by siRNA further protected *Raver1^−/−^* immortalized macrophages from TNF/LPS+TAK1-i-induced cytotoxicity (Figures 5B and S6D). Furthermore, the inclusion of the *Ripk1* alternative exon is further increased in this condition, compared to either deletion of *Raver1* or knockdown of *Ptbp1* (Figure 5C). Consistently, the level of canonically spliced *Ripk1* mRNA is reduced (Figure 5D) upon targeting both splicing factors, resulting in lower protein levels of full-length RIPK1 than *Raver1^−/−^* macrophages or *Ptbp1* knockdown alone (Figure 5E). Together, these data suggest that Raver1 cooperates with Ptbp1 to regulate *Ripk1* pre-mRNA splicing and cell death signaling.

### Identification of additional innate immune signals regulated by Raver1

We performed a broad transcriptome analysis to explore if Raver1 regulates the expression of genes other than RIPK1. RNA-seq data indicated that *Ripk1* read counts were reduced, and the expression of only a small number of genes was influenced by the absence of *Raver1* in macrophages (Figures 6A and S7A). The majority of the innate immunity pathways tested appeared largely preserved in *Raver1*^−/−^ BMDMs (Figure S7A). For example, only small, non-significant differences were observed in LPS– and TNF-induced NF-κB activation (Figures S7B-S7D). Notable exceptions were polyI:C-TLR3 (Figures S7E and S7F) and ssRNA encephalomyocarditis virus (EMCV)-MDA5 (Figure S7F); responses to both these stimulants were reduced in *Raver1*^−/−^ BMDMs. This is consistent with a potential contribution of RIPK1, as previous work demonstrates that TLR3 drives the association of RIPK1 with TRIF (38). Prior work has also implicated Raver1 in the regulation of MDA5 signaling (39), which is triggered by EMCV (40).

**Figure 6.**
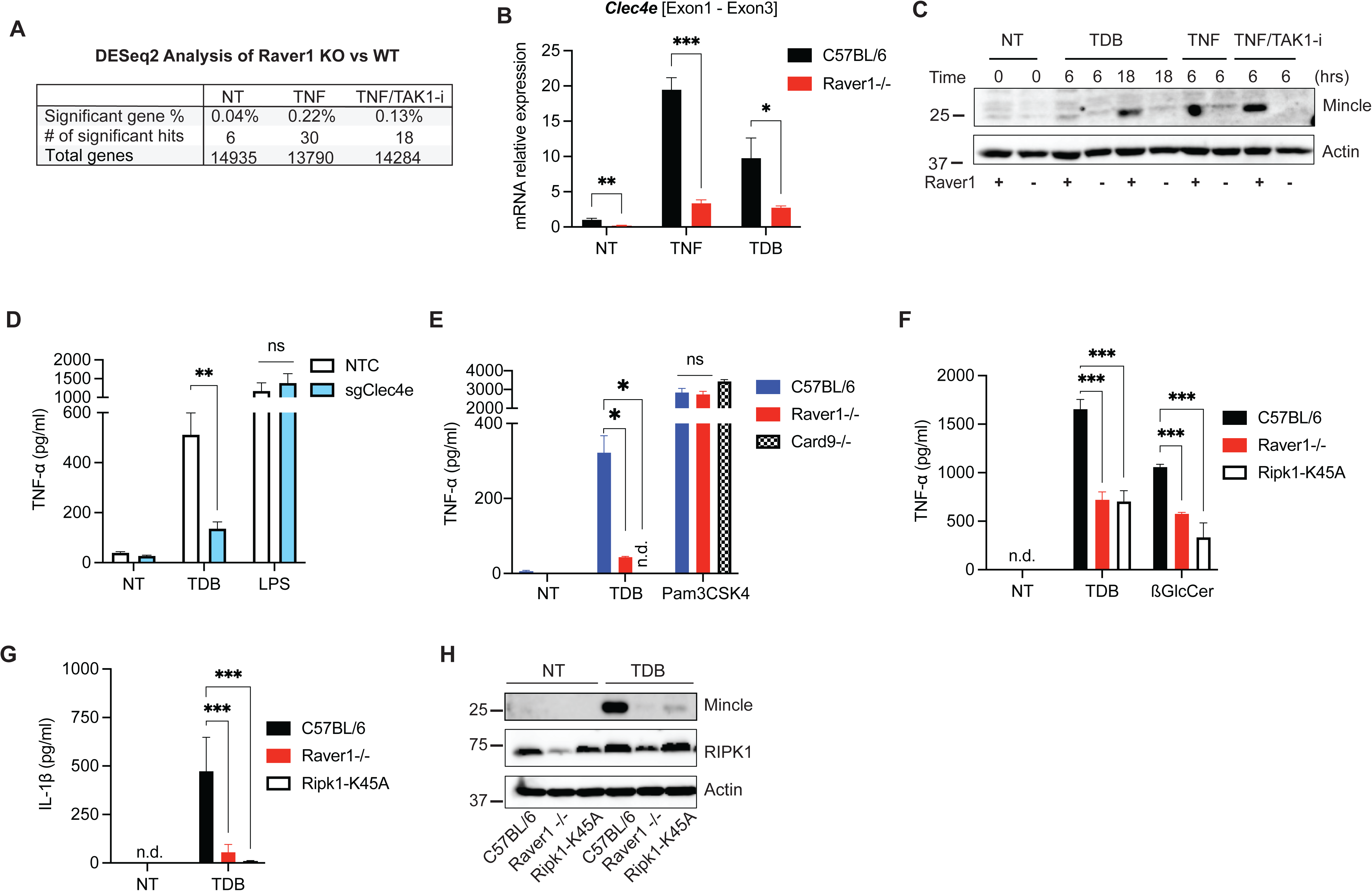
Raver1 modulates Mincle-dependent TNF release via RIPK1. (**A**) WT or *Raver1*^−/−^ BMDMs were unstimulated (NT) or stimulated for 1 hr with TNF or TNF plus TAK1-i and analyzed by RNAseq. Genes with significant differences in expression between WT and *Raver1*^−/−^ are enumerated. (**B, C**) WT or *Raver1*^−/−^ BMDMs were stimulated with TNF or synthetic mycobacterial cord factor, trehalose-6,6’-dibehenate (TDB). (B) Relative mRNA expression of Mincle (*Clec4e*) was quantified by qRT-PCR after 1-hour TNF or 6-hour TDB treatment, normalized to *Actin*. (C) Protein lysates were analyzed for Mincle level by immunoblotting at the indicated times. (**D-G**) BMDMs transduced with *Clec4e* sgRNA (D) or from indicated genotypes (E-G) were stimulated with TDB, LPS, Pam_3_CSK_4_, or cell-death-related β-glucosylceramide (βGlcCer) for 24 hours. TNF-α or IL-1β release was assessed by ELISA. (**H**) WT, *Raver1^−/−^*, and *Ripk1^K45A^* BMDMs were treated with TDB for 18 hours and then analyzed by immunoblotting. Data are presented as the mean ± SD for triplicate wells from three or more independent experiments. Immunoblots are representative of three blots performed. For comparison between two groups, Student’s t-test was used; for more than two groups, ANOVA was used with Bonferroni’s post-hoc test. n.s., not significant; *p≤0.05; **p≤0.01; ***p≤0.001. *Abbreviations*: NTC, non-targeting control; n.d., not detected.

RNA-seq analysis also identified macrophage *Cd4* mRNA as being downregulated and potentially influenced by Raver1 (Figure S7A); however, further analysis revealed no significant differences in CD4 protein expression levels and CD4^+^ T cell composition in spleens from *Raver1*^−/−^ mice (Figure S7G and S7H). The composition of other spleen lymphoid and myeloid populations was also similar in the absence of *Raver1* (Figure S7H), indicating that immune cell composition is largely preserved in *Raver1*^−/−^ animals.

### Raver1 regulates Mincle-dependent TNF release via RIPK1

One of the few genes with reduced baseline gene expression in the absence of *Raver1* (Figures S7A) was *Clec4e* (encoding Mincle). Mincle is a C-type lectin receptor (CLR) that plays a pivotal role in recognizing various pathogens and damaged cells. It initiates NF-κB activation via CARD9 and is susceptible to upregulation by TNF (41, 42). Subsequent analysis indicated a sharp reduction in Mincle-*Clec4e* mRNA (Figure 6B) and protein expression (Figure 6C), as well as cellular responses in *Raver1*^−/−^ BMDMs to two Mincle ligands: synthetic mycobacterial cord factor, trehalose-6,6’-dibehenate (TDB) (43) and cell-death-associated ß-glucosylceramide (ßGlcCer) (44) (Figures 6D-6G). However, we did not observe significant differences in alternative splicing events for *Clec4e* in the absence of *Raver1* (Figures S8A and S8B); thus, the contribution of Raver1 on Mincle-*Clec4e* gene expression could be indirect. One connection between Mincle and Raver1 could be linked to the capacity of RIPK1 to modulate CLR signaling through CARD9 (45). Notably, using RIPK1-K45A kinase-dead mutant BMDMs, we found a marked reduction in TDB-induced TNF and IL-1ß transcription and release (Figures 6F, 6G, and S8C). Consistently, TDB-induced Mincle expression is substantially reduced in the absence of RIPK1 kinase activity (Figure 6H). Therefore, we propose that Raver1 modulation of RIPK1 impacts Mincle signaling.

### Raver1 is indispensable for host defense against *Yersinia* infection

To investigate the biological significance of Raver1 in host defenses *in vivo*, we infected *Raver1*^−/−^ mice orally with *Y. pseudotuberculosis*. Caspase-8 and RIPK1 kinase play central roles in host resistance to this gastroenteric pathogen (Figures S9A and S9B) (13, 14, 46). Compared to C57Bl/6 (*Raver1*^+/+^) or *Raver1*^+/-^ littermates, *Raver1*^−/−^ mice had markedly elevated susceptibility to bacterial challenge (Figure 7A), although the gross composition of immune cells in the spleens was preserved (Figure S9C). Compared to WT mice, bacterial loads in the spleens and livers of *Raver1*^−/−^ mice increased significantly (Figure 7B). Our experiments with BMDMs suggested that Raver1 has a major effect on *Yersinia*-induced cell death (Figure 2). Consistent with this, we also observed a significant decrease in cytotoxicity in monocytes and macrophages isolated from infected *Raver1*^−/−^ mice (Figures 7C and S9D). Histological analysis of liver tissue showed that increased bacterial loads correlated with increased number and size of inflammatory foci (Figures 7D and 7E). Notably, visible bacterial microcolonies were observed only in *Raver1*^−/−^ but not in C57Bl/6 mice (Figures 7F and 7G), indicating that *Raver1*^−/−^ mice had a diminished ability to contain bacterial spread and replication. Therefore, recruited inflammatory cells in *Raver1*^−/−^ livers may not display an appropriate antibacterial capacity, as we have proposed by observing livers from mice with other defects in innate immune signaling (29). Moreover, IL-18 cytokine levels were markedly downregulated in spleens of *Raver1^−/−^*mice (Figure 7H), although IL-1ß, TNF, and IL-6 were not significantly changed (Figures S9E-S9G). IL-18 is a critical cytokine for host resistance to *Yersinia* (29, 31, 47). Specifically, we found that genetic ablation of *Il18* resulted in heightened susceptibility to *Y. pseudotuberculosis* gastrointestinal infection (Figure 7I), following increased bacteria load in spleens (Figure 7J), emphasizing the importance of normal IL-18 production in host defenses to this pathogen. Our results indicate that Raver1 regulates *Yersinia*-induced IL-18 production both *in vivo* (Figure 7K) and in macrophages *in vitro* (Figure 3J), and the evidence indicates that the Raver1-RIPK1-caspase-8-IL-18 axis is a key contributor to host defenses. We also noticed a small but significant decrease in the percentage of CD69^+^ B and CD4^+^ T cell populations in *Yersinia*-infected *Raver1*^−/−^ mice (Figures S9H-S9J), suggesting that Raver1 contributes to the development of adaptive immune responses. This is likely a consequence of positive regulation of early innate immune responses to bacteria, by aiding lymphoid cell activation. Collectively, these findings indicate that Raver1 expression is critical for resistance to infection and to coordinate an appropriate host response to *Yersinia* bacteria.

**Figure 7.**
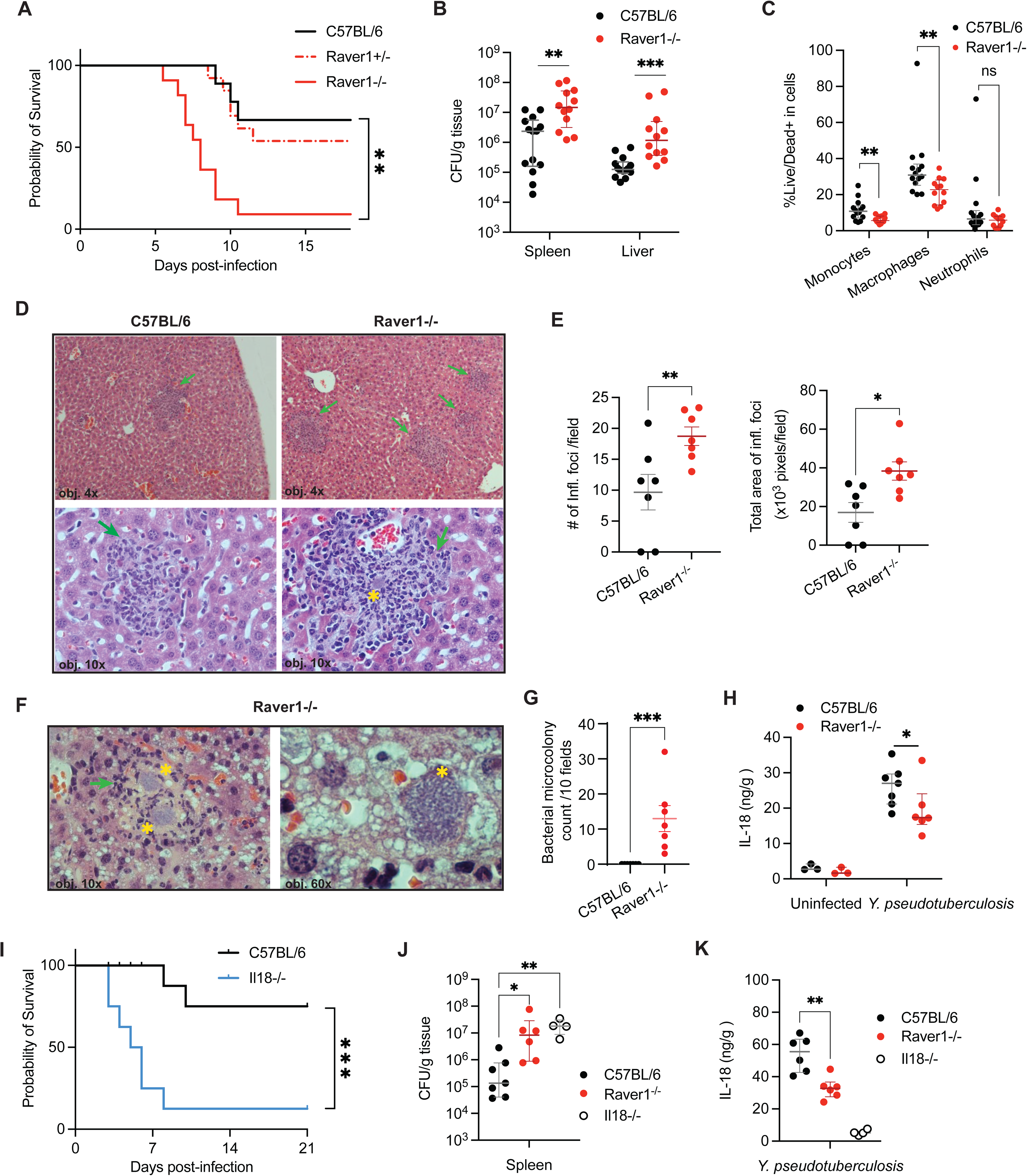
Raver1 is essential for host resistance against *Y. pseudotuberculosis* infection *in vivo*. (**A**) Survival of C57BL/6 (n=9), *Raver1*^−/−^ (n=11), or heterozygous littermates (n=13) were monitored following oral challenge with 0.5–1 x 10^8^ CFU of *Y. pseudotuberculosis.* (**B-C**) tissues from C57BL/6 (n=14) and *Raver1*^−/−^ (n=12) mice were collected five days post-infection to quantify the bacterial load in spleens and livers (B), or to analyze subpopulations of CD11b^+^ myeloid cells in spleens for cell death with live/dead stain by flow cytometry (C). (**D-G**) Liver sections from C57BL/6 (n=7) and *Raver1*^−/−^ (n=7) mice were stained with hematoxylin and eosin and subjected to microscopy. (E) Inflammatory foci (green arrows) and (G) bacterial microcolonies (yellow asterisks) were quantified. Magnification: 40x (4x objective, as indicated), 100x (10x objective), or 600x (60x objective). (**H**) IL-18 in C57BL/6 (n=7) and *Raver1*^−/−^ (n=6) mice spleen homogenates three days post-infection were measured by ELISA. (**I**) Survival of C57BL/6 (n=8) and *Il18^−/−^*(n=8) mice challenged orally with 1 x 10^8^ CFU of *Y. pseudotuberculosis.* (**J**) Bacterial loads in indicated spleens were quantified three days post-infection. (**K**) IL-18 in indicated spleen homogenates were measured by ELISA three days post-infection. Data are presented as medians with interquartile range (B, C, H, J, K) or mean with SEM (E, G), either pooled from two independent experiments (A-C) or representative of two or more independent experiments (D-K). Images are representative of 2 individual experiments. For comparisons, data sets were analyzed by log-rank test (A, I), or Mann-Whitney U test (B, C, E, G, H, K), Kruskal-Wallis with Dunn’s post-hoc test (J). *p≤0.05, **p≤0.01, ***p≤0.001.

## Discussion

Alternative splicing plays a crucial role in modulating inflammation and cell death responses by influencing isoform expression of key pathway factors. Here, we identified Raver1 as a crucial upstream regulator of RIPK1-caspase-8-mediated pyroptosis and IL-1ß/IL-18 secretion, uncovering a new regulatory mechanism in the interplay between immune responses and cell death pathways. Our data support a role for Raver1 in maintaining RIPK1 abundance and function by repressing the inclusion of an alternative exon during pre-mRNA splicing. These results provide the first evidence of Raver1 in the regulation of innate immunity and cell death, in primary macrophages and *in vivo* (Figure S10).

Our data suggest that Raver1 is a pivotal regulator, orchestrating multiple interconnected pathways. In addition to a profound effect on pyroptotic/apoptotic cell death and inflammatory responses (IL-18/IL-1ß release) to *Yersinia* bacteria and the type III secretion system effector molecule YopJ, we also found that Raver1 positively influenced caspase-8-mediated signaling induced by LPS, TNF family members, and IFNγ, and restrained by kinases TAK1 and IKK (Figures 1–3). In contrast, canonical caspase-1-dependent pyroptosis and IL-1ß/IL-18 release, and raptinal-induced intrinsic apoptosis were unaffected by Raver1. Moreover, Raver1 facilitates MLKL-dependent necroptosis unleashed by caspase-8 blockade, influencing the plasticity between cell death pathways through RIPK1 (48, 49).

Raver1 is ubiquitously expressed in cells and tissues, and its promoter region contains a negative regulator element that could be influenced by NF-κB (50). This supports our observations of decreased *Raver1* transcription upon TLR4 stimulation (Figure 1K) and the presence of a proposed negative feedback loop. Raver1 impacts cell death and inflammation across various cell types, including macrophages, neutrophils, osteosarcoma, and fibroblast cells, suggesting that Raver1 may play a fundamental role in coordinating immune responses throughout the body in pathological conditions via its regulation of RIPK1. Additionally, the regulatory role of Raver1 spans species, effectively governing cell death processes in both mouse and human cells. This versatile function underscores its potential significance as a key regulator in cellular homeostasis and immune responses.

In this study, we determined alternative splicing as an important layer of regulation on RIPK1 function in addition to well-established post-translational modifications. Alternative splicing can profoundly impact the balance between cell survival and death. By altering splicing patterns, cells can modulate the expression levels or activities of cell death or survival factors.

For example, alternative splicing can generate splice variants that either promote or inhibit apoptosis. These variants may possess different protein domains. For example, different splice variants of *cflar* (cFlip) and *bcl-x* can either induce or protect cells from cell death (12, 51, 52). The RIPK1 Splice I variant is incapable of inducing cell death by itself, as it lacks a significant portion of the RIPK1 protein (Figure 1G). Interestingly, we found Splice I directly interacted with full-length RIPK1 and attenuated its activation in a dominant-negative manner, potentially by blocking phosphorylation at S166 (Figure 4). Mechanistically, we speculate that Splice I binding to RIPK1 restrains RIPK1 kinase-dependent recruitment of FADD and caspase-8, thus impeding complex II assembly, which is limited in *Raver1*^−/−^ macrophages (Figure 2J). Our data revealed reduced newly synthesized but not pre-existing RIPK1 protein in the presence of Splice I. One possible explanation is that the absence of *Raver1* and altered splicing could mediate the mRNA decay of steady-state *Ripk1*. Together, the reduction in full-length RIPK1 expression along with the dominant-negative effect of RIPK1 non-functional variant Splice I, explain the decrease of pyroptosis/apoptosis in *Raver1*^−/−^ cells during TAK1 inhibition.

Raver1 has been proposed as a regulatory factor of Ptbp1, yet not many Raver1 targets have been identified. While Ptbp1 has a significant impact on a wide range of splicing events from regulation of neuronal differentiation to lymphocyte maturation and activation (22, 53–56), the specific contribution of Raver1 to innate immune function remains largely unexplored. Here, we conclude that signaling induced by both pathogens and cytokines, in primary macrophages and mice, is modulated by the action of Raver1. This indicates a previously unknown and broad impact on innate immune responses, in part due to alternative splicing regulation. Transcriptomic analyses revealed that loss of Raver1 resulted in few perturbations to mRNA levels, but a pervasive and widespread effect on transcript composition (Figure 1). We show here that Raver1 influences mRNA isoform diversity primarily through repression of alternative exon inclusion.

We also demonstrated that regulation of transcript composition in targets, such as *Ripk1*, has direct effects on physiologically relevant innate immune processes. Further analysis could identify additional immune-related target transcripts of Raver1 and the potential involvement of Ptbp1. These two splicing factors can function together to regulate the splicing of certain genes (19, 20), while Ptbp1-independent alternative splicing by Raver1 is also possible (37). Our data suggest that Raver1 may work in concert with Ptbp1 in regulating *Ripk1* pre-mRNA splicing and subsequent cytotoxicity, as evidenced by the additive effect of modulating Raver1 and Ptbp1 (Figure 5). Additional studies are needed to more precisely define the interplay between Ptbp1 and Raver1 during innate immune signaling. It is tempting to speculate that Raver1 provides further specificity to the action of Ptbp1. Raver1 was proposed to bridge multiple Ptbp1 molecules bound to polypyrimidine tracts flanking exons to promote RNA looping and splicing repression (21). One possible model is that Raver1 serves as a connection between Ptbp1 proteins flanking the *Ripk1* alternative exon to collaboratively promote its skipping.

Interestingly, we found the expression and function of the CLR Mincle (*Clec4e*) were also impaired in *Raver1*^−/−^ macrophages, although we did not detect a direct link to the splicing of *Clec4e* (Figure 6). Here we provide evidence that Raver1 controls Mincle via RIPK1 kinase activity. Notably, Dectin-1 (*Clec7a*) has been proposed to relay necroptosis signals via RIPK1, and an interaction with RIPK1 and the CARD9 protein, which also contributes to Mincle signaling, was proposed in macrophages (45). It is possible that Mincle-driven CARD9 serves as an adaptor for RIPK1 binding and activation, controlling downstream NF-κB-activated transcription of pro-inflammatory cytokines and Mincle (*Clec4e*). This could potentially serve as a novel mechanism with a feed-forward loop to enhance immunity against infection.

Here we demonstrate the pathophysiological significance of proper *Ripk1* splicing by Raver1 in the context of *Yersinia* challenge (Figure 7). *Raver1* deficiency results in decreased inflammatory cytokine release, reduced cell death of host phagocytes, and increased bacterial propagation, culminating in decreased host survival. Surprisingly, we observed a significant decrease in splenic IL-18 but not IL-1ß. This finding may be explained by cytokine upregulation influenced by intensified bacterial burden, and the impact of different caspases contributing to the maturation of IL-18 and IL-1ß under different experimental conditions (57). In this context, a recent study proposed that IL-1ß is less important for mouse survival against *Y. pseudotuberculosis* infection (58), whereas our experiments show that IL-18 is a strong positive factor in host defenses. Our findings indicate that Raver1 may not primarily influence the recruitment of phagocytes to infected sites, but rather is essential for their effective anti-bacterial functions. In addition, we noted a significant proportional decrease in activated CD69^+^ spleen T and B cells upon deletion of *Raver1*, indicative of a potential impact on adaptive immune responses. Regulation of IL-18 production may also influence adaptive immune responses. Raver1 may contribute to lymphocyte priming through IL-18 regulation, thereby assisting adaptive immunity.

*Yersinia* infections serve as an excellent model to study the RIPK1-caspase-8-GSDMD axis with potential implications in different immune contexts. RIPK1 contributes to many disease conditions, including bacterial infection (46), dermatitis (59), autoinflammatory diseases (60, 61), cancer (3) and neuroinflammation (62). Additionally, a link between caspase-8 and GSDMD has been proposed for intestinal inflammation triggered by mutation of FADD (63).

These intersecting crosstalk pathways are prime targets for drug development. Decreased expression of TAK1 in the aging process has also been implicated in neuroinflammation (64), and TAK1 inhibition is being tested for cancer therapy (65, 66). Thus, our signaling studies are relevant for pathological conditions beyond bacterial infection. Interestingly, a recent study identified three cases of patients carrying homozygous mutations in RIPK1 causing recurrent infections and severe inflammatory dysregulation (67). These mutations resulted in premature stop codon within RIPK1 kinase domain, producing truncated proteins, much like Splice I in Raver1-deficient conditions. Moreover, genome-wide association studies have suggested that SNPs within the *RAVER1* locus are associated with COVID-19 disease severity and also inflammatory diseases (68, 69), indicating that regulation of Raver1 could impact both infectious and sterile inflammatory conditions. Taken together, our data provide a better understanding of this new layer of splicing regulation and could hold promise in aiding development of therapeutic strategies to control and fine-tune inflammatory responses.

## Materials and Methods

### Mice

All experiments involving mice were approved by the University of Massachusetts Chan Medical School’s Institutional Animal Care and Use Committee. Most mouse strains used in this study were described previously (13, 15, 29, 30) and bred in-house. *Raver1*^−/−^ mice were generated in-house with the UMass Chan Transgenic Mouse Core using CRISPR/Cas9 technology. Briefly, *Cas9* mRNA together with *in vitro* transcribed sgRNA targeting exon 1 of *Raver1* (GCCGCAAGATACTGATCCG) was injected into mouse zygotes to generate *Raver1*^−/−^ mice. Detailed mouse strain information as described in SI Materials and Methods.

### Bacterial strains

*Y. pestis* (KIM5), *Y. pseudotuberculosis* IP32777, *Y. enterocolitica* JA127/90, their derivative strains, and *Salmonella enterica* serovar Typhimurium strain SL1344, were used as previously described (13, 15, 31). Detailed bacterial strain information and growth condition described in SI Materials and Methods.

### Cell culture

Bone marrow-derived macrophages (BMDMs) were differentiated from bone marrow harvested from femurs and tibia of 6–20-week-old mice, in DMEM supplemented with 10% fetal calf serum (FCS), 25 mM HEPES, and 10% L929 conditioned medium containing M-CSF for 6 days, as described (30, 31). Detailed information as described in SI Materials and Methods.

### CRISPR cell death screens

Forward genetic genome-wide screens were performed using the mouse BRIE knockout CRISPR pooled library designed by David Root and John Doench (70) (Addgene 73633) in Cas9-expressing BMDMs. Detailed information as described in SI Materials and Methods.

### Data availability

All data needed to evaluate the conclusions in this paper are present either in the main text or in the supplementary materials. MLKL KO mice were obtained via a material transfer agreement (MTA) with the Walter and Eliza Hall Institute of Medical Research. Caspase 8-deficient mice were obtained via an MTA with the University Health Network, Toronto. RIPK3 KO mice, GSDMD KO mice, and mouse GSDMD antibody were obtained via an MTA with Genentech. RIPK1 kinase-dead KI K45A mice and RIPK3 and RIPK1 inhibitors were obtained via an MTA with GlaxoSmithKline.

## Acknowledgments

This manuscript is dedicated to the memory of our colleague Jon D. Goguen. He passed away in July 2022 and contributed to this study with bacterial strains and discussions thereof. The UMass Chan Medical School Transgenic Animal Modeling Core is acknowledged for assistance with generation of *Raver1*^−/−^ animals. We thank C. Choquette and Z. Jiang for help with mice, M. Trombly for assistance with manuscript development, V. Boyartchuk for discussions, E. Pearlman for advice on neutrophil experiments, and A. Olive, C. Sassetti, P. Meraner, and A. Brass for input on CRISPR screen methods.

## Author contributions

The authors’ contributions are as follows, Conceptualization: EL, KAF, PO; Reagents: JB, MKP, LR, TE; Investigation: BZ, PO, AD, RE; Discussions, analyses: JL, LTU, AAP, MAK, RK, TE; Funding acquisition: EL, KAF, AAP, MAK, TE, RK; Supervision: EL, KAF, AAP; Writing – original draft: PO, BZ, EL; Writing – review and editing: all authors.

---

**Figure S1. CRISPR screen identifies Raver1 as a regulator controlling RIPK1-induced cell death.** (**A**) Cas9-BMDMs transduced with indicated sgRNAs were stimulated with *Y. pestis* or LPS plus TAK1 inhibitor (TAK1-i) and cell death was measured by LDH release after 4 hours. (**B**) sgRNA design and *Raver1* DNA sequence in generating *Raver1*^−/−^ mouse strains #4 and #6. Strain #6 is used as *Raver1*^−/−^ throughout the study, except where noted. (**C**) Cell lysates from C57BL/6, *Raver1*^−/−^ #4 or #6 BMDMs were analyzed for Raver1 expression, confirming knockout. (**D**) Sashimi plot of the Raver1 gene from RNA-seq of C57BL/6 or Raver1 BMDMs. (**E**) Ridge plot showing directionality of changes in alternative splicing events from RNA-seq data in *Raver1^−/−^ vs.* WT BMDMs. Delta percent spliced-in (PSI) of *Raver1^−/−^* vs. WT BMDMs is shown on the x-axis, with alternative splicing event types shown on the y-axis. For comparison, two-sided t-tests were used between significant and non-significant deltaPSI of each alternative splicing event type; ns, not significant; *p≤0.05, ***p≤0.001. (**F**) RNA-seq read coverage (RPKM, y-axis) over *Ripk1* exons 4 and 5 (genomic coordinates on chr13 are shown on the x-axis) for untreated *Raver1* WT (left) and *Raver1*^−/−^ (right) BMDMs. Reads are summed across three independent replicates. Curved lines show positions of exon-exon junction reads in the region and junction read counts are in red text. The y-axis scale is adjusted to account for the differential gene expression of *Ripk1* between *Raver1*-WT and *Raver1*^−/−^ mice. (**G**) Relative mRNA levels of *Ripk1* fragments in BMDMs of indicated genotypes with or without *Y. pestis* infection are shown. Quantification of expression of exon4-alternative exon and exon4-exon5 was analyzed by qRT-PCR. (**H, I**) Cell lysates from Cas9-expressing BMDMs transduced with *Raver1*-specific sgRNA or *Raver1^−/−^* BMDMs from 5 mice were analyzed by immunoblotting. (I) Samples from the same experiment as Fig. 1J are shown. Data are presented as the mean ± SD for triplicates from three or more independent experiments. Immunoblots are representative of 3 immunoblots performed. For comparison, ANOVA with Bonferroni’s post-hoc test was used for (A), Student’s t-test was used for (G). n.s., not significant; *p≤0.05; **p≤0.01, ***p≤0.001. *Abbreviations*: n.d., not detected; NTC, non-targeting control; A3SS, alternative 3′ splice sites; A5SS, alternative 5′ splice sites; MXE, mutually exclusive exons; RI, retained intron; SE, skipped exons; Alt exon, alternative exon.

**Figure S2. Raver1 mediates RIPK1-caspase-8-dependent cell death and IL-1β/IL-18 cytokine release.** (**A–E**) Indicated BMDMs (A–C, E) or peritoneal macrophages (D) were infected with *Y. pestis* or challenged with TAK1-i plus LPS, TNF, or LT-α. Cell death was measured by the percentage of SYTOX-positive cells over time (A, D, E) or LDH release after 4 hours (B, C). BMDMs in (C) were derived from mouse strain #4. (**F**) Caspase-8 enzymatic activity was measured by Caspase-8-Glo in WT or *Raver1^−/−^* BMDMs 2 hours after stimulation with indicated reagents. (**G, H**) BMDM cell lysates were analyzed by immunoblotting following stimulation with indicated ligands for the indicated times (G) or following infection with *Y. pestis* for 2 hours (H). Data are presented as the mean ± SD for triplicate wells from three or more independent experiments. Immunoblots are representative of 3 immunoblots performed. For comparison between two groups, Student’s t-test was used; for more than two groups, ANOVA with Bonferroni’s post-hoc test was used. n.s., not significant; *p≤0.05, **p≤0.01, ***p≤0.001.

**Figure S3. Raver1 controls IL-1β/IL-18 release and inflammation in response to *Yersinia* bacterial infection.** (**A, B**) BMDMs (A) or neutrophils (B) were infected or challenged for 5 hours with indicated ligands. Supernatants were used for ELISA to measure the release of IL-1β. (**C**) Combined cell extracts and supernatants were analyzed by immunoblotting 3 hours after *Y. pestis* infection on BMDMs. (**D**) Oligomerization of ASC in the inflammasome-enriched and crosslinked lysates was detected by immunoblotting in BMDMs treated with TAK1-i plus LPS or TNF for 3 hours. (**E, F**) BMDMs primed with Pam_3_CSK_4_ (1 μg/ml) for 3 hours were infected with *Salmonella*, *Y. pestis*, *Y. pestis* mutant (*Y. pestis* ΔYopMΔYopJ), or challenged with ATP (5 mM), Nigericin (10 μM), TcdB (50 μg/ml), poly[dA:dT] (4 μg/ml), TAK1-i plus TNF, or TAK1-i plus LT-α. IL-1β was measured by ELISA after 5 hours (E). Cell death was measured by LDH release after 4 hours (F). Data are presented as the mean ± SD for triplicate wells from three or more independent experiments. Immunoblots are representative of 3 immunoblots performed. For comparison between two groups, Student’s t-test was used; for more than two groups, ANOVA with Bonferroni’s post-hoc test was used. n.s., not significant; *p≤0.05, **p≤0.01, ***p≤0.001. n.d., not detected.

**Figure S4. Raver1 promotes IFNγ-triggered cell death dependent on RIPK1, caspase-8, and GSDMD.** (**A, B**) C57BL6 BMDMs were treated with IFNγ, TAK1-i, or both for the indicated times. Cell death was measured by LDH release (A) or SYTOX dye membrane penetration (B). (**C, D**) BMDMs (C) or peritoneal macrophages (D) of indicated genotypes were stimulated with IFNγ or IFNγ+TAK1-i. Cell death was measured by LDH release after 4.5 hours (C), or SYTOX dye membrane permeability (D). (**E, F**) Indicated BMDMs were challenged with IFNγ+TAK1-i for 3 hours. Caspase-8 activity was measured by Caspase-8-Glo (E). Cleavage of GSDME and caspase-3 was detected by immunoblotting (F). (**G**) Indicated BMDMs were treated with IFNγ+LPS or IFNγ+TAK1-i. Cell death was measured by SYTOX dye membrane penetration. Data are presented as the mean ± SD for triplicate wells from three or more independent experiments. Immunoblots are representative of 3 immunoblots performed. For comparison between two groups, Student’s t-test was used; for more than two groups, ANOVA was used with Bonferroni’s post-hoc test. n.s., not significant; ***p≤0.001.

**Figure S5. Raver1 dysfunction results in a RIPK1 splicing isoform, Splice I, that negatively regulates cell death.** (**A**–**C**) Cas9-expressing WEHI164 clone 13 cells (A) or human U2OS cells (B, C) were challenged with TRAIL, TNF, or TNF plus TAK1-i, with or without a 1-hour pretreatment with pan-caspase inhibitor zVAD-fmk (25 μM), RIP1[i] (1 μM), or RIP3[i] (3 μM). Cell death was measured by SYTOX dye membrane permeability (A) or LDH release after 10 hours (B, C). (**D**) Doxycycline-inducible vectors containing RIPK1 Splice I were transfected or transduced into HEK293T or WEHI164 clone 13 cells. Doxycycline (Dox) was added for the indicated times. Cell lysates were analyzed for full-length RIPK1 or HA-Splice I by immunoblotting. Data are presented as the mean ± SD for triplicate wells from three or more independent experiments. Immunoblots are representative of 3 immunoblots performed. For comparison, ANOVA was used with Bonferroni’s post-hoc test. n.s., not significant; *p≤0.05; **p≤0.01; ***p≤0.001. *Abbreviations*: RIP1[i], RIPK1 inhibitor; RIP3[i], RIPK3 inhibitor; hTNF, human TNF.

**Figure S6. Raver1 cooperates with Ptbp1 to regulate RIPK1 splicing and cell death**. (**A, B**) mRNA levels from resting Cas9-expressing BMDMs transduced with sgRNA against *Raver1* or *Ptbp1* were analyzed. (A) Visualization of PCR fragments of *Ripk1* exon 4 to exon 5 showed increased fragment size (184 bp vs. 73 bp), suggesting the inclusion of the alternative exon (Alt. Exon). (B) Quantification of exon4-alternative exon vs. exon4-exon5 was performed by qRT-PCR. (**C, D**) WT or *Raver1^−/−^* immortalized macrophages were transfected with Ptbp1-targeting siRNA or non-targeting control (NTC) siRNA. *Ptbp1* mRNA levels were quantified by qRT-PCR to evaluate knockdown efficiency (C). Cell death was measured by SYTOX membrane permeability following TNF+TAK1-i treatment (D). *Ptbp1* knockdown by siRNA displayed a similar but weaker phenotype than *Raver1*-deficient macrophages, potentially due to compensation by Ptbp2, a functional redundant protein for Ptbp1. (B, C) mRNA levels were normalized to *Actin*. Data are presented as the mean ± SD for triplicates representative of 3 independent experiments. For comparison between two groups, Student’s t-test was used; for more than two groups, ANOVA was used with Bonferroni’s post-hoc test. n.s., not significant; ***p≤0.001.

**Figure S7. Raver1 regulates additional innate immune signals**. (**A**) Heatmap from RNAseq, showing a small set of genes with significant changes in mRNA levels in *Raver1*^−/−^(KO) compared to *Raver1*^+/+^ (WT) BMDMs that were untreated (NT) or incubated with TNF. Genes with transcription level of log_2_ fold-change >0.6 and adjusted P-value (padj) <0.01 are considered significant (★). Specifically, the transcription of related immune and cell death pathway genes is largely preserved in *Raver1*^−/−^. (**B-D**) *Raver1* WT or knockout BMDMs were stimulated with LPS or TNF for 1 hour (B) or as indicated (C, D). Cell lysates were quantified for *Il1b* and *Tnf* relative mRNA levels by qRT-PCR (B, normalized to *Actin*), or analyzed for NF-κB signaling protein processing by immunoblotting (C, D). (**E, F**) The TNF-α production from BMDMs treated with stated stimuli at 24 hours was detected by ELISA. (E) Pam_3_CSK_4_ (TLR2 ligand, 1 μg/ml), poly(I:C) (TLR3 ligand, 20 μg/ml), LPS (TLR4 ligand, 100 ng/ml), or R848 (TLR7/8 ligand, 1 μg/ml) was directly added to cells; 5’-ppp-dsRNA (RIG-I ligand, 1 μg/ml) and 2’3’-cGAMP (cGAS/STING ligand, 1 μg/ml) were transfected into cells after 1-hour priming with LPS (10 ng/ml). (F) Macrophages were infected with EMCV at MOI=10 for MDA5 pathway activation. (**G, H**) Immune cell composition in uninfected spleens of *Raver1*^−/−^ and WT mice identified by flow cytometry. (G) Percentage of CD4^+^ splenocytes in CD3^+^ T cells are shown. (H) The data indicate that immune cell composition is similar in the presence or absence of *Raver1*. Group size: n=3 for *Raver1*^−/−^ and C57BL/6. Data are presented as the mean ± SD (B-F) or median with 95% interquartile range (H) for triplicates representative of 2–3 independent experiments. For comparison between two groups, Student’s t-test was used; for more than two groups, ANOVA was used with Bonferroni’s post-hoc test; Mann-Whitney U test was used for (H). n.s., not significant; *p≤0.05; **p≤0.01; ***p≤0.001.

**Figure S8. Raver1 modulates Mincle-dependent TNF release via RIPK1.** (**A**) Relative expression of Mincle (*Clec4e*) fragment in nontreated (NT) or TNF-treated WT or *Raver1*^−/−^ BMDMs were measured by qRT-PCR. (**B**) Boxplots of Mincle (*Clec4e*) alternative splicing as detected by rMATS run on RNA-seq data from *Raver1^−/−^* or WT BMDMs, with percent spliced-in (PSI) on the y-axis, and alternative splicing event (ASE) types on the x-axis. No events were detected for alternative 3′ splice sites (A3SS), alternative 5′ splice sites (A5SS), mutually exclusive exons (MXE), and retained intron (RI). SE, skipped exons. (**C**) WT, *Raver1^−/−^* and *Ripk1^K45A^* BMDMs were treated with TDB for 6 hours. *Il1b* relative mRNA levels were quantified by qRT-PCR. (A, C) mRNA levels were normalized to *Actin*. Data are presented as the mean ± SD for triplicates representative of 3 independent experiments. For comparison between two groups, Student’s t-test was used; for more than two groups, ANOVA was used with Bonferroni’s post-hoc test. n.s., not significant; *p≤0.05; **p≤0.01; ***p≤0.001.

**Figure S9. Raver1, RIPK1 kinase activity, and caspase-8 play essential roles in host resistance against *Y. pseudotuberculosis* infection**. (**A–B**) Mice with indicated genotypes were infected with 0.5–1 x 10^8^ CFU of *Y. pseudotuberculosis* by oral gavage. Survival was monitored for up to 28 days. (**C–J**) C57BL/6 or *Raver1*^−/−^ mice were challenged orally with 0.5–1 x 10^8^ CFU of *Y. pseudotuberculosis.* Tissues were collected 3 days (E-G) or 5 days (C, D, H–J) post-infection. Immune cell composition in spleen was detected by flow cytometry in (C). Subpopulations of CD11b^+^ myeloid cells in spleens were analyzed for cell death with live/dead stain by flow cytometry (D). IL-1β, TNF, and IL-6 in spleen homogenates were measured by ELISA (E-G). (H–J) Flow cytometry was performed to detect lymphocyte activation by CD69 antibodies. Percentage and median fluorescence intensity (MFI) of CD69^+^ cells are shown. Data are presented as median with interquartile range (C, E–I) representative of two or three independent experiments. Group size: (A) C57Bl/6, n=6; *Ripk1^K45A^*, n=6; *Ripk1^D138N^*, n=7; (B) C57Bl/6, n=6; *Mlkl*^−/−^*Casp8*^+/-^, n=6; *Mlkl*^−/−^*Casp8*^−/−^, n=5; (C-J) C57Bl/6, n=7, *Raver1*^−/−^, n=6 (E-G) or n=7 (C, D, H-J). For comparison, data sets were analyzed by log-rank Mantel-Cox tests (A, B) and Mann-Whitney U tests (C–J). n.s., not significant; *p≤0.05.

**Figure S10. Proposed model for Raver1-mediated caspase-8-dependent pyroptotic cell death, inflammation, and pathogen resistance.** In the presence of Raver1 (left), RIPK1 abundance is maintained by Raver1 canonical splicing. During infection, TLR or TNF signaling together with TAK1 or IKK inhibition by effector protein YopJ lead to activation of caspase-8 via RIPK1. Active caspase-8 initiates caspase-3-dependent apoptosis, GSDMD-dependent pyroptosis, and promotes IL-1β and IL-18 maturation, facilitating cytokine release. This enables proper inflammation in mice to limit bacterial replication, thereby resisting pathogen invasion. In the absence of Raver1 (right), RIPK1 is alternatively spliced, causing decreased full-length RIPK1 and increased splice variant (Splice I). Splice I interacts with RIPK1 and inhibits its function. The reduced overall RIPK1 level together with the dominant negative function of Splice I, cooperatively result in compromised cell death and inflammation in *Raver1^−/−^*mice, culminating in susceptibility to bacterial infection.

## References

1. B. Tummers, D. R. Green, Caspase-8: regulating life and death. Immunol Rev 277, 76–89 (2017).

2. P. Orning, E. Lien, Multiple roles of caspase-8 in cell death, inflammation, and innate immunity. J Leukoc Biol 109, 121–141 (2021).

3. J. Clucas, P. Meier, Roles of RIPK1 as a stress sentinel coordinating cell survival and immunogenic cell death. Nat Rev Mol Cell Biol 24, 835–852 (2023).

4. N. Paquette et al., Serine/threonine acetylation of TGFbeta-activated kinase (TAK1) by Yersinia pestis YopJ inhibits innate immune signaling. Proc Natl Acad Sci U S A 109, 12710–12715 (2012).

5. U. Meinzer et al., Yersinia pseudotuberculosis effector YopJ subverts the Nod2/RICK/TAK1 pathway and activates caspase-1 to induce intestinal barrier dysfunction. Cell Host Microbe 11, 337–351 (2012).

6. S. Mukherjee et al., Yersinia YopJ acetylates and inhibits kinase activation by blocking phosphorylation. Science 312, 1211–1214 (2006).

7. R. Mittal, S. Y. Peak-Chew, H. T. McMahon, Acetylation of MEK2 and I kappa B kinase (IKK) activation loop residues by YopJ inhibits signaling. Proc Natl Acad Sci U S A 103, 18574–18579 (2006).

8. R. K. S. Malireddi et al., TAK1 restricts spontaneous NLRP3 activation and cell death to control myeloid proliferation. J Exp Med 215, 1023–1034 (2018).

9. K. Maeda, J. Nakayama, S. Taki, H. Sanjo, TAK1 Limits Death Receptor Fas-Induced Proinflammatory Cell Death in Macrophages. J Immunol 209, 1173–1179 (2022).

10. Y. Dondelinger et al., NF-kappaB-Independent Role of IKKalpha/IKKbeta in Preventing RIPK1 Kinase-Dependent Apoptotic and Necroptotic Cell Death during TNF Signaling. Mol Cell 60, 63–76 (2015).

11. Y. Dondelinger et al., Serine 25 phosphorylation inhibits RIPK1 kinase-dependent cell death in models of infection and inflammation. Nat Commun 10, 1729 (2019).

12. H. I. Muendlein et al., cFLIP(L) protects macrophages from LPS-induced pyroptosis via inhibition of complex II formation. Science 367, 1379–1384 (2020).

13. D. Weng et al., Caspase-8 and RIP kinases regulate bacteria-induced innate immune responses and cell death. Proc Natl Acad Sci U S A 111, 7391–7396 (2014).

14. N. H. Philip et al., Caspase-8 mediates caspase-1 processing and innate immune defense in response to bacterial blockade of NF-kappaB and MAPK signaling. Proc Natl Acad Sci U S A 111, 7385–7390 (2014).

15. P. Orning et al., Pathogen blockade of TAK1 triggers caspase-8-dependent cleavage of gasdermin D and cell death. Science 362, 1064–1069 (2018).

16. S. B. Kovacs, E. A. Miao, Gasdermins: Effectors of Pyroptosis. Trends Cell Biol 27, 673–684 (2017).

17. J. Sarhan et al., Caspase-8 induces cleavage of gasdermin D to elicit pyroptosis during Yersinia infection. Proc Natl Acad Sci U S A 115, E10888–E10897 (2018).

18. Z. Zheng et al., The Lysosomal Rag-Ragulator Complex Licenses RIPK1 and Caspase-8-mediated Pyroptosis by Yersinia. Science 372 (2021).

19. S. Huttelmaier et al., Raver1, a dual compartment protein, is a ligand for PTB/hnRNPI and microfilament attachment proteins. J Cell Biol 155, 775–786 (2001).

20. N. Gromak et al., The PTB interacting protein raver1 regulates alpha-tropomyosin alternative splicing. EMBO J 22, 6356–6364 (2003).

21. A. P. Rideau et al., A peptide motif in Raver1 mediates splicing repression by interaction with the PTB RRM2 domain. Nat Struct Mol Biol 13, 839–848 (2006).

22. N. Keppetipola, S. Sharma, Q. Li, D. L. Black, Neuronal regulation of pre-mRNA splicing by polypyrimidine tract binding proteins, PTBP1 and PTBP2. Crit Rev Biochem Mol Biol 47, 360–378 (2012).

23. Y. Zhao et al., The NLRC4 inflammasome receptors for bacterial flagellin and type III secretion apparatus. Nature 477, 596–600 (2011).

24. M. G. Callow et al., CRISPR whole-genome screening identifies new necroptosis regulators and RIPK1 alternative splicing. Cell Death Dis 9, 261 (2018).

25. R. K. S. Malireddi et al., Whole-genome CRISPR screen identifies RAVER1 as a key regulator of RIPK1-mediated inflammatory cell death, PANoptosis. iScience 26, 106938 (2023).

26. K. W. Chen et al., RIPK1 activates distinct gasdermins in macrophages and neutrophils upon pathogen blockade of innate immune signaling. Proc Natl Acad Sci U S A 118 (2021).

27. L. Laurien et al., Autophosphorylation at serine 166 regulates RIP kinase 1-mediated cell death and inflammation. Nat Commun 11, 1747 (2020).

28. A. Polykratis et al., Cutting edge: RIPK1 Kinase inactive mice are viable and protected from TNF-induced necroptosis in vivo. J Immunol 193, 1539–1543 (2014).

29. G. I. Vladimer et al., The NLRP12 inflammasome recognizes Yersinia pestis. Immunity 37, 96–107 (2012).

30. D. Ratner et al., The Yersinia pestis Effector YopM Inhibits Pyrin Inflammasome Activation. PLoS Pathog 12, e1006035 (2016).

31. D. Ratner et al., Manipulation of Interleukin-1beta and Interleukin-18 Production by Yersinia pestis Effectors YopJ and YopM and Redundant Impact on Virulence. J Biol Chem 291, 9894–9905 (2016).

32. Y. Zheng et al., A Yersinia effector with enhanced inhibitory activity on the NF-kappaB pathway activates the NLRP3/ASC/caspase-1 inflammasome in macrophages. PLoS Pathog 7, e1002026 (2011).

33. F. R. Greten et al., NF-kappaB is a negative regulator of IL-1beta secretion as revealed by genetic and pharmacological inhibition of IKKbeta. Cell 130, 918–931 (2007).

34. R. Karki et al., Synergism of TNF-alpha and IFN-gamma Triggers Inflammatory Cell Death, Tissue Damage, and Mortality in SARS-CoV-2 Infection and Cytokine Shock Syndromes. Cell 184, 149–168 e117 (2021).

35. D. S. Simpson et al., Interferon-gamma primes macrophages for pathogen ligand-induced killing via a caspase-8 and mitochondrial cell death pathway. Immunity 55, 423–441 e429 (2022).

36. T. Espevik, J. Nissen-Meyer, A highly sensitive cell line, WEHI 164 clone 13, for measuring cytotoxic factor/tumor necrosis factor from human monocytes. J Immunol Methods 95, 99–105 (1986).

37. A. Wedler et al., RAVER1 hinders lethal EMT and modulates miR/RISC activity by the control of alternative splicing. Nucleic Acids Res 10.1093/nar/gkae046 (2024).

38. E. Meylan et al., RIP1 is an essential mediator of Toll-like receptor 3-induced NF-kappa B activation. Nat Immunol 5, 503–507 (2004).

39. H. Chen et al., RAVER1 is a coactivator of MDA5-mediated cellular antiviral response. J Mol Cell Biol 5, 111–119 (2013).

40. L. Gitlin et al., Essential role of mda-5 in type I IFN responses to polyriboinosinic:polyribocytidylic acid and encephalomyocarditis picornavirus. Proc Natl Acad Sci U S A 103, 8459–8464 (2006).

41. J. Schick et al., Cutting Edge: TNF Is Essential for Mycobacteria-Induced MINCLE Expression, Macrophage Activation, and Th17 Adjuvanticity. J Immunol 205, 323–328 (2020).

42. E. C. Patin, S. J. Orr, U. E. Schaible, Macrophage Inducible C-Type Lectin As a Multifunctional Player in Immunity. Front Immunol 8, 861 (2017).

43. H. Schoenen et al., Cutting edge: Mincle is essential for recognition and adjuvanticity of the mycobacterial cord factor and its synthetic analog trehalose-dibehenate. J Immunol 184, 2756–2760 (2010).

44. M. Nagata et al., Intracellular metabolite beta-glucosylceramide is an endogenous Mincle ligand possessing immunostimulatory activity. Proc Natl Acad Sci U S A 114, E3285–E3294 (2017).

45. M. Cao et al., Dectin-1-induced RIPK1 and RIPK3 activation protects host against Candida albicans infection. Cell Death Differ 26, 2622–2636 (2019).

46. L. W. Peterson et al., RIPK1-dependent apoptosis bypasses pathogen blockade of innate signaling to promote immune defense. J Exp Med 214, 3171–3182 (2017).

47. E. Bohn et al., IL-18 (IFN-gamma-inducing factor) regulates early cytokine production in, and promotes resolution of, bacterial infection in mice. J Immunol 160, 299–307 (1998).

48. K. Newton et al., Activity of caspase-8 determines plasticity between cell death pathways. Nature 575, 679–682 (2019).

49. M. Fritsch et al., Caspase-8 is the molecular switch for apoptosis, necroptosis and pyroptosis. Nature 575, 683–687 (2019).

50. M. G. Romanelli, P. Lorenzi, F. Avesani, C. Morandi, Functional characterization of the ribonucleoprotein, PTB-binding 1/Raver1 promoter region. Gene 405, 79–87 (2007).

51. D. R. Ram et al., Balance between short and long isoforms of cFLIP regulates Fas-mediated apoptosis in vivo. Proc Natl Acad Sci U S A 113, 1606–1611 (2016).

52. C. F. A. Warren, M. W. Wong-Brown, N. A. Bowden, BCL-2 family isoforms in apoptosis and cancer. Cell Death Dis 10, 177 (2019).

53. E. Monzon-Casanova et al., The RNA-binding protein PTBP1 is necessary for B cell selection in germinal centers. Nat Immunol 19, 267–278 (2018).

54. J. La Porta, R. Matus-Nicodemos, A. Valentin-Acevedo, L. R. Covey, The RNA-Binding Protein, Polypyrimidine Tract-Binding Protein 1 (PTBP1) Is a Key Regulator of CD4 T Cell Activation. PLoS One 11, e0158708 (2016).

55. S. Vavassori, Y. Shi, C. C. Chen, Y. Ron, L. R. Covey, In vivo post-transcriptional regulation of CD154 in mouse CD4+ T cells. Eur J Immunol 39, 2224–2232 (2009).

56. J. A. Hensel et al., Splice factor polypyrimidine tract-binding protein 1 (Ptbp1) primes endothelial inflammation in atherogenic disturbed flow conditions. Proc Natl Acad Sci U S A 119, e2122227119 (2022).

57. B. Demarco et al., Caspase-8-dependent gasdermin D cleavage promotes antimicrobial defense but confers susceptibility to TNF-induced lethality. Sci Adv 6 (2020).

58. R. Matsuda et al., A TNF-IL-1 circuit controls Yersinia within intestinal pyogranulomas. J Exp Med 221 (2024).

59. S. B. Berger et al., Cutting Edge: RIP1 kinase activity is dispensable for normal development but is a key regulator of inflammation in SHARPIN-deficient mice. J Immunol 192, 5476–5480 (2014).

60. N. Lalaoui et al., Mutations that prevent caspase cleavage of RIPK1 cause autoinflammatory disease. Nature 577, 103–108 (2020).

61. P. Tao et al., A dominant autoinflammatory disease caused by non-cleavable variants of RIPK1. Nature 577, 109–114 (2020).

62. L. Mifflin, D. Ofengeim, J. Yuan, Receptor-interacting protein kinase 1 (RIPK1) as a therapeutic target. Nat Rev Drug Discov 19, 553–571 (2020).

63. R. Schwarzer, H. Jiao, L. Wachsmuth, A. Tresch, M. Pasparakis, FADD and Caspase-8 Regulate Gut Homeostasis and Inflammation by Controlling MLKL– and GSDMD-Mediated Death of Intestinal Epithelial Cells. Immunity 52, 978–993 e976 (2020).

64. D. Xu et al., TBK1 Suppresses RIPK1-Driven Apoptosis and Inflammation during Development and in Aging. Cell 174, 1477–1491 e1419 (2018).

65. R. Santoro, C. Carbone, G. Piro, P. J. Chiao, D. Melisi, TAK-ing aim at chemoresistance: The emerging role of MAP3K7 as a target for cancer therapy. Drug Resist Updat 33-35, 36–42 (2017).

66. A. Singh et al., TAK1 inhibition promotes apoptosis in KRAS-dependent colon cancers. Cell 148, 639–650 (2012).

67. D. Cuchet-Lourenco et al., Biallelic RIPK1 mutations in humans cause severe immunodeficiency, arthritis, and intestinal inflammation. Science 361, 810–813 (2018).

68. D. Ellinghaus et al., Analysis of five chronic inflammatory diseases identifies 27 new associations and highlights disease-specific patterns at shared loci. Nat Genet 48, 510–518 (2016).

69. I. M. Fink-Baldauf, W. D. Stuart, J. J. Brewington, M. Guo, Y. Maeda, CRISPRi links COVID-19 GWAS loci to LZTFL1 and RAVER1. EBioMedicine 75, 103806 (2022).

70. J. G. Doench et al., Optimized sgRNA design to maximize activity and minimize off-target effects of CRISPR-Cas9. Nat Biotechnol 34, 184–191 (2016).

