## Supplementary Information for "Raver1 links *Ripk1* RNA splicing to caspase-8-mediated pyroptotic cell death, inflammation, and pathogen resistance"

This PDF file includes:

- SI Materials and Methods
- Figures S1 to S10
- Tables S1 and S2
- SI References

### SI Materials and Methods

Mice

All experiments involving mice were approved by the University of Massachusetts Chan Medical School's Institutional Animal Care and Use Committee. Most mouse strains used in this study were described previously (1-4) and bred in-house. *Rip3*$^{-/-}$*Casp8*$^{-/-}$ rescued the embryonic lethality of *Casp8*$^{-/-}$ mice (5), and were originally bred by W. Kaiser and E. Mocarski. *Rip3*$^{-/-}$ mice (6) were generated by K. Newton and V. Dixit (Genentech) and provided by Drs. W. Kaiser, E. Mocarski, C. Dillon, and D. Green. *Gsdmd*$^{-/-}$ mice (7) were generated at Genentech by V. Dixit, B. Lee, and N. Kayagaki. *Ripk1*$^{D138N}$ mutant mice (8) were from M. Kelliher. *Ripk1*$^{K45A}$ kinase-dead mutant mice (9) were generated at GlaxoSmithKline (GSK) and provided by J. Bertin. *Gsdme*$^{-/-}$ mice were purchased from Jax (#032411) (10). *Mlkl*$^{-/-}$ mice were provided by F. Chan and originally obtained from W. Alexander (11). *Ifngr1*$^{-/-}$ mice (Jax #3288) (12) were obtained from J. Harris, and *Tnfr1*$^{-/-}$ mice were from Jax (#3242) (13). *Card9*$^{-/-}$ mice were provided by C. Specht and S. Levitz (Jax #28652) (14). *Il18*$^{-/-}$ mice were provided by D. Golenbock (Jax #004130). Transgenic *Cas9* mice expressing a 3x-FLAG tagged *Streptococcus pyogenes Cas9*, linked to EGFP via a self-cleaving peptide P2A were from Jax (#26179) (15). For CRISPR cell death screens, Cas9-expressing *Rip3*$^{-/-}$ cells were used. *Raver1*$^{-/-}$ mice were generated in-house with the UMass Chan Transgenic Mouse Core using CRISPR/Cas9 technology. Briefly, *Cas9* mRNA together with *in vitro* transcribed sgRNA targeting exon 1 of *Raver1* (GCCGCAAGATACTGATCCG) was injected into mouse zygotes to generate *Raver1*$^{-/-}$ mice. Two separate genetically defined mouse strains were studied (strains #6 and #4), and BMDMs from the two mouse strains behaved similarly in terms of decreased responses to stimulatory conditions triggering RIPK1-caspase-8. All data were generated from mouse strain #6 except for data shown in Figure S2C, which are from mouse strain #4.

Antibodies and reagents

Antibodies used in this study include RIPK1 (38/RIP(RUO), BD), phospho-RIPK1 (S166, CST), caspase-1 (#AG-20B-0042-C100, Adipogen), ASC (D2W8U, CST), GSDMD (17G2G9, Genentech) (16), GSDME (EPR19859, Abcam), IL1β (AF-401-NA, R&D), cleaved caspase-8 (D5B2, CST), full-length caspase-8 (1G12, Enzo), cleaved caspase-3 (5A1E, CST), phospho-IKKα/β (Ser176/180) (16A6, CST), IκBα (CST, 9242), phospho-p65 (93H1, CST), FLAG (R&D, A8592), HA (Sigma, H6908), Raver1 (MBS9203524, MyBioSource), Mincle (D292-3, MBL) and Actin (A3854, Sigma). Reagents used in this study include: Pam3CSK4 (Invivogen), repurified TLR2 ligand-free LPS (*E. coli* O111:B4, Sigma) (17), R848 (Invivogen), and poly(I:C) (Invivogen). We also used recombinant mouse TNF-α (Peprotech), recombinant mouse lymphotoxin-alpha (LT-α) (BIOMATIK), recombinant mouse IFNγ (Peprotech), recombinant mouse TRAIL (Peprotech), and recombinant human TNF-α (Peprotech). Furthermore, we used RIPK1 inhibitor (GSK'963, GlaxoSmithKline or Selleckchem) (1, 18), RIPK3 inhibitor (GSK'872, GlaxoSmithKline or Selleckchem) (19), pan-caspase inhibitor zVAD-fmk (Millipore), TAK1 inhibitor (TAK1-i) 5z-7-oxozeaneol (abbreviated to 5z7, EMD), and IKK inhibitor (TPCA-1, Tocris Bioscience) (20). In addition, we used cycloheximide (CHX) and raptinal, a stimulator of intrinsic apoptosis (Adipogen). ATP (Sigma Aldrich) and Nigericin (Invivogen) were purchased from commercial sources. Some experiments included pyrin-activating *Clostridioides* (previously *Clostridium)* toxin TcdB (a gift from Dana Borden Lacy) (21), poly(dA:dT) (Invivogen, tlrl-patn), cGAMP (Invivogen), and 5'ppp-dsRNA (Invivogen). The encephalomyocarditis virus (EMCV) used in experiments was strain VR-129B (ATCC) (22). Cells were also exposed to a synthetic version of mycobacterial cord factor, trehalose-6,6'-

dibehenate TDB (Invivogen), or cell death-related β-glucosylceramide (βGlcCer) (Invivogen). The SYTOX Orange used for kinetic cell death measurements was purchased from Invitrogen and Hoechst dye from ThermoFisher. The DSS crosslinker used for ASC oligomerization assays was purchased from ThermoFisher. For lactate dehydrogenase (LDH) release cell death assays, we used the CytoTox 96 kit (Promega). Reduced ATP production in dying cells was measured by CellTiter-Glo (Promega). To determine caspase-8 enzymatic activity towards a tetra-peptide substrate, we used Caspase-8 Glo (Promega). Cytokine release was measured by ELISA using IL-1β, TNF, and IL-6 kits from R&D Systems, and IL-18 from Invitrogen.

Bacterial strains and growth conditions

The pgm-deficient type III secretion system (T3SS) pCD1+ strain of *Y. pestis* (KIM5), as well as its isogenic T3SS mutant derivatives (ΔYopJ, ΔYopMΔYopJ), were described previously (2, 3, 17). The *Y. pseudotuberculosis* IP32777 (serogroup O:1, also known as IP2777) and IP32777ΔYopJ (23-25), were provided by I. Brodsky and J. Bliska. *Y. enterocolitica* JA127/90 (serogroup O:8 biotype IA – a wild-type, virulent strain) and A127/90 ΔYopP (called YopJ in *Y. pestis* and *Y. pseudotuberculosis*) were provided by R. Adkins and G. Cornelis (26). *Y. pestis* and *Y. pseudotuberculosis* strains were grown in tryptose-beef extract (TB) broth with 2.5 mM $CaCl_2$, and *Y. enterocolitica* strains were grown in LB broth. All *Yersinia* strains were plated overnight from frozen glycerol stocks and then grown at 26 ˚C in liquid broth overnight; on the day of infection, cultures were diluted 1:10 and grown for 1 hour at 26 ˚C followed by a shift to 37 ˚C for 2 hours. Bacteria were then washed three times in PBS, quantified by OD600, and added to cells at a multiplicity of infection (MOI) of 10 (*Y. pestis*, *Y. enterocolitica*) or 40 (*Y. pseudotuberculosis*) CFU per cell. *Salmonella enterica* serovar Typhimurium strain SL1344 (provided by M. O'Riordan), also expressing T3SS, was grown in LB medium at 37 ˚C. *Salmonella* bacteria were added to cells at an MOI of 2. Gentamicin (50 μg/ml) was added to cultures 2.5 h post-infection to limit extracellular replication of *Yersinia* and *Salmonella* bacteria, except for experiments shown in Figure 2D, in which case gentamicin (8 μg/ml) was added 30 min post-infection.

Cell culture

Bone marrow-derived macrophages (BMDMs) were differentiated from bone marrow harvested from femurs and tibia of 6–20-week-old mice, in DMEM supplemented with 10% fetal calf serum (FCS), 25 mM HEPES, and 10% L929 conditioned medium containing M-CSF for 6 days, as described (2, 27). Neutrophils were elicited by intraperitoneal injection of 1 ml of 9% casein (Sigma Aldrich) 16 and 3 hours prior to peritoneal cavity lavage. Neutrophils were isolated from peritoneal fluid and enriched using the EasySep Mouse negative selection kit (StemCell) (28). Neutrophil purity was over 95%, as determined by flow cytometry gating on $CD45^+CD11b^+Ly6G^+Ly6C^{lo}$. Cells were plated in RPMI with 10% FBS, 1% non-essential amino acids (Gibco), 1% sodium pyruvate (Gibco), and 10 ng/mL granulocyte-macrophage colony-stimulating factor (Peprotech) for 3 hours before the experiment. Peritoneal macrophages were recruited by injection of 2 ml thioglycolate (Remel) in mice 3 days before lavage. For cell lines, immortalized WT and *Raver1*$^{-/-}$ macrophages (Figure 6) were generated from C57BL/6 and *Raver1*$^{-/-}$ BMDMs through transduction of J2 retrovirus expressing c-myc and c-raf. Immortalized macrophages, mouse fibrosarcoma WEHI164 clone 13 cells (29), and human osteosarcoma U2OS cells (30, 31) were cultured in complete DMEM.

Cell stimulations

Cells were stimulated with bacterial strains as described above, or with the chemical inhibitors 5z7 TAK1-i (0.4 μM), TPCA-1 IKK-i (5 μM), cycloheximide (CHX, 2 μg/ml), raptinal (10 μM), SMAC mimetic Birinapant (5 μM), GSK'963 RIPK1-i (1 μM), GSK'872 RIPK3-i (3 μM), and zVAD pan-caspase-inhibitor (25 μM). For GSK'963, GSK'872, and zVAD, these inhibitors were added 1 h before the addition of subsequent stimuli. Concentrations of stimuli were: LPS (100 ng/ml), TNF (mouse, 10 ng/ml), LT-α (10 ng/ml), IFNγ (50 ng/ml), TRAIL (50 ng/ml), hTNF (10 ng/ml), unless otherwise stated. For RIPK1-caspase-8 pathway stimulation, LPS, TNF family cytokines, or IFNγ were added to cells simultaneously as TAK1-i or IKK-i. For inflammasome activation, macrophages were primed with LPS or Pam3CSK4 (1 μg/ml) for 3 hours before stimulation with nigericin (10 μM), ATP (5 mM), TcdB (50 ng/ml), or transfected with poly(dA:dT) (4 μg/ml). All IL-1β and IL-18 experiments were performed for 5 h. For NF-κB pathway activation, macrophages were treated with the TLR ligand Pam3CSK4 (1 μg/ml), poly(I:C) (20 μg/ml), LPS (100 ng/ml), or R848 (1 μg/ml). 5'-ppp-dsRNA (1 μg/ml), and 2'3'-cGAMP (1 μg/ml) were transfected into cells after 1-hour priming with LPS (10 ng/ml). For MDA5 pathway activation, macrophages were infected with EMCV (MOI=10) (32). After 24 hours, supernatant was collected for TNF ELISA. For Mincle stimulation, cells were treated with trehalose-6,6'-dibehenate (TDB) (10 μg/ml), or plate-coated βGlcCer (75 μg/million cells). Supernatant was collected for TNF or IL-1β ELISA after 24 hours, cell lysates were collected for RNA after 6 hours, or protein after 18 hours.

Cell death assays

The LDH assay (CytoTox 96, Promega) was used to measure cell death at a 4-hr endpoint, unless otherwise stated in the figure. For this assay, the medium was replaced with phenol red-free DMEM supplemented with 3.5% FCS and 25 mM HEPES prior to infection. CellTiter-Glo (Promega) was also used for end-point cell death, measuring ATP. For this experiment, complete DMEM was used. Additionally, a kinetic cell death assay using a DNA-binding dye was performed as follows. The cells were incubated with 200 nM SYTOX Orange dye (Invitrogen) and Hoechst (ThermoFisher) in the DMEM medium mentioned above right after adding stimuli. Upon stimulation, the plate was placed in a Cytation5 Imager (BIOTEK) at 37 ˚C with 5% $CO_2$, and fluorescence measurements were made every 15 min. Increased fluorescence correlates with DNA binding by SYTOX Orange upon entry through increasingly permeable cell membranes as cell death progresses. Hoechst was used as a counterstain for quantification of total cell counts.

Immunoblots and immunoprecipitations

Cells were harvested at different time points in 1xLDS buffer (ThermoFisher) with 5 mM DTT, protease and phosphatase inhibitor cocktails (Roche). For caspase-1 and IL-1β immunoblot, a combined cell lysate and supernatant was loaded on the gel. For other blots, only the cell lysate was loaded on SDS gel. For immunoprecipitations, cells were washed once and lysed in Triton buffer for 30 minutes (1% Triton X-100, 150 mM NaCl, 5 mM KCl, 2 mM $MgCl_2$, 1 mM EDTA, 25 mM Tris-HCl, pH 7.4) with protease and phosphatase inhibitor. Lysates were cleared by centrifugation at 12,000*g* and a fraction was saved as input control. The remaining lysate was cleared once with protein G agarose beads (ThermoFisher) to remove non-specific protein binding, followed by incubation with pre-blocked beads for 30 min, then the addition of a pull-down antibody against caspase-8 or RIPK1 overnight at 4 ˚C. Beads were then washed 4 times in Triton buffer, and bound proteins were eluted by direct addition of SDS loading buffer with 5 mM DTT. The immunoprecipitated proteins and saved lysates were analyzed by immunoblot.

ASC oligomerization

Cross-linking assays of higher-order ASC protein complexes were performed as previously described (33). Briefly, experimentally treated cells were harvested, washed, and lysed in NP-40 lysis buffer. After removing cell debris, the supernatant was centrifuged at a series of high speeds to separate large protein complexes. The lysate was kept as an input control, and the pellet was washed with CHAPS buffer (0.1% CHAPS, 150 mM KCl, 50 mM HEPES, pH 7.4) and cross-linked with DSS. Both fractions were then resuspended in SDS loading buffer and analyzed by immunoblot.

Quantitative real-time PCR

RNA was extracted using the RNeasy (Qiagen) or Aurum (BioRad) RNA extraction kits. RNA was then transformed into cDNA using the iScript cDNA Synthesis kit (BioRad). cDNA was mixed with iTaq Universal SYBR Green Supermix (BioRad) and F/R primers. PCR products were monitored in triplicate on a BioRad CFX96. Relative gene expression was determined using the $2^{-ddCt}$ method normalized to the housekeeping genes *Actin* or *Gapdh*. Primers used for qPCR are shown in Table S1.

CRISPR

C57BL/6 BMDMs endogenously expressing Cas9 (15) and Cas9-expressing WEHI164 clone13 cells were transduced with lentivirus containing lentiGuide-Puro cloned with sgRNA using the F. Zhang protocol (34). sgRNA sequences were chosen using the Doench algorithm (35) and the online web tool Benchling (Table S2). sgRNAs cloned into the lentiGuide-Puro vector (Addgene 52963) were packaged into lentivirus using plasmids psPAX2 (Addgene 12260) and pMD2 (Addgene 12259). Briefly, 4 million HEK293T cells were plated in a 10-cm dish one day prior to transfection. Cells were then transfected using lipofectamine with 6 µg lentiGuide-Puro, 4 µg psPAX2, and 2 µg pMD2 following the manufacturer's protocol. 48 hours post-transfection, virus-containing supernatant was collected and filtered through a 0.45 µm PVDF filter.

CRISPR cell death screens

Forward genetic genome-wide screens were performed using the mouse BRIE knockout CRISPR pooled library designed by David Root and John Doench (36) (Addgene 73633). Briefly, four sgRNAs targeting every coding gene in the mouse genome as well as 1000 non-targeting controls were packaged into lentivirus using HEK293T cells and transduced into primary Cas9-expressing BMDMs. Cells were transduced at ~1000x coverage of the BRIE library using a low multiplicity of infection (MOI<0.3) as follows. Bone marrow from five *Rip3*$^{-/-}$ mice expressing Cas9 (15) were plated in 10-cm non-TC treated plates in DMEM supplemented with 10% FBS and 20% L929 medium. At two days post isolation, cells were transduced with the BRIE pooled library. At day three, the medium was replaced with fresh DMEM medium containing FBS and L929. At day four, 5 µg/ml puromycin was added to the cells. At day five, another 10 ml of fresh medium containing L929 and 5 µg/ml puromycin was added to the cells. At day seven, mature, transduced, BMDMs were harvested, counted, and ~80 million cells were plated in four 15 cm TC dishes in medium without antibiotics. The following day, cells were challenged with *Y. pestis* KIM5 at an MOI of 10 for 5 hours. Cells were then harvested, washed in 1x PBS, and stained with live/dead blue stain (Invitrogen). Cells were then fixed in 2% PFA for 20 minutes and the live cell population was sorted on a BD FACS Aria cell sorter. The live cell population was lysed, and DNA was harvested using a Dneasy Blood and Tissue kit (Qiagen).

Libraries for sequencing were prepared as previously described (36). After PCR amplification of isolated genomic DNA with Illumina compatible primers, PCR products were purified using

AMPure DNA-binding beads and sequenced on an Illumina NextSeq 500. Sequence reads were trimmed to adjust for p5 primer stagger using cutadapt (v2.9). Reads were then analyzed using MAGeCKFlute (37).

RNA-seq library prep

RNA-seq samples were prepared from WT or Raver1 KO BMDMs from three biological replicates (cells from three different mice) seeded in 6-well plates at 2 million cells per well. The cells were stimulated with TNF (10 ng/ml) or TNF+TAK1-inhibitor (0.4 µM) for 1 hour. RNA was extracted using the RNeasy RNA extraction kit (Qiagen). A pair-end RNA-seq library was generated using the TruSeq Stranded Total RNA Library Prep Gold Kit (Illumina). Samples were then quality-controlled using FragmentAnalysis and sequenced on a NovaSeq S4 flowcell.

RNA-seq analysis

FASTQ files were analyzed using the DolphinNext Platform developed by Yukselen et al. (38). Briefly, sequencing reads were mapped to mouse_mm10_refseq and quantified using Kallisto. Differential expression analysis was performed using DESeq2 with NT vs TNF or TNF/TAK1-i, or NT vs TNF + TNF/TAK1-i. Significant hits were selected based on a cutoff of padj of 0.01 and a Log2(fold change) of 0.6.

RNA-seq coverage plots were made using "bedtools coverage" to get read counts for all basepairs in the *Ripk1* genes and counts were normalized and plotted in R. RNA-seq read coverage in reads per kilobase per million (RPKM), positions of exon-exon junction reads in the region and junction read counts are shown. Reads are summed across three independent replicates.

Splicing analysis

For alternative splicing analyses, high-throughput sequencing reads were mapped with STAR v2.7.0 (39) (default parameters) to the GRCm38.p6 (mm10) reference genome. Alternative splicing was quantified with rMATS v4.1.0 (40), scaffolded on the RefSeq mm10 annotations (downloaded from UCSG Genome Browser), with parameters "—libType fr-firststrand --readLength 151 -variable-read-length." Alternative splicing changes were assessed by comparing *Raver1*$^{-/-}$ replicates and wild-type (WT) replicates. Splicing events with FDR ≤ 10% were identified as significant.

Splice variant studies

For interaction studies, pcDNA3.1-HA-RIPK1 was co-transfected with pcDNA3.1 cloned with FLAG-Splice I, FLAG-kinase domain (KD), or empty vector into 5 million HEK293T cells. Cells were lysed for immunoprecipitation or immunoblotting 24 hours after transfection. For cell death assays, lentivirus containing doxycycline-inducible vector pTRIPz-RIPK1-HA (a gift from C. Park and F. Chan) or Splice I-HA were transduced into HEK293T or WEHI164 clone 13 cells and antibiotics-selected for 3 days. 1–2 µg/ml Doxycycline (Sigma) was used to induce protein expression. For some experiments, after 12 hours of doxycycline treatment, TNF (50 ng/ml) was added to WEHI164 clone 13 cells, pcDNA3.1-HA-RIPK1 or empty vector was transfected into HEK293T cells to measure cell death.

DsiRNA transfection

TriFECTa® RNAi Kit was ordered from IDT. Predesigned DsiRNA duplexes for mouse *Ptbp1* were used (mm.Ri.Ptbp1.13.1, mm.Ri.Ptbp1.13.2, mm.Ri.Ptbp1.13.3), along with a non-targeting control (NTC). Immortalized macrophages derived from *Raver1*$^{+/+}$ (WT) or *Raver1*$^{-/-}$ mouse were subjected to DsiRNA transfection using Lipofectamine® RNAiMAX Reagent following manufacturer's instructions (Invitrogen). Cells were rested for 48-72 hours before stimulations or lysis for protein or RNA extractions.

*In vivo* infection and tissue harvest

6–12-week-old mice were challenged with *Y. pseudotuberculosis* (IP32777) via oral gavage using a dose of 0.5–1 x$10^8$ colony forming units (CFUs)/mouse. Bacteria were prepared after TB plate growth as mentioned above. Survival was monitored every 12 hours. For organ harvest experiments, mice were euthanized by $CO_2$ at 3 days or 5 days post-infection. Spleen and liver (one lobe per mouse) were harvested and homogenized in gentleMACS™ C Tubes. A portion was taken out and plated at 10-fold dilutions on LB or TB plates for bacterial load count (CFU/g tissue). For spleens, supernatants from the homogenate were used for cytokine measurement by ELISA, and cells from the pellet were used for flow cytometry. Liver pathology is very instructive in terms of understanding tissue inflammation during *Yersinia* infections (1, 3, 17, 27). One lobe of the liver was fixed in 4% paraformaldehyde overnight and stained with hematoxylin and eosin for histology. Sections were imaged using a Nikon TE400 microscope with a DS-Fi1 camera. Inflammatory foci number and size were traced manually through ImageJ by taking the average in 3 microscopic fields/sections, excluding foci size less than 500 pixel$^2$ (pixels squared). Visible bacterial microcolonies were quantified by the average of 10 microscopic fields/section. Two sections were sliced per mouse.

Flow cytometry

To measure cell death and activation of different splenocyte populations, single-cell suspensions were plated in a 96-well plate at 1 million cells/well and stained with Live/Dead Blue (Invitrogen L34962) for viability, following the manufacturer's instructions. Cells were Fc-receptor blocked (2.4G2, Tonbo) then stained with the following antibodies: CD45 (30-F11), CD11c (HL3), CD11b (M1-70), Ly6C (AL-21), CD4 (GK1.5), and CD8a (53-6.7) (all from BD Biosciences), as well as Ly6G (1A8) and CD3 (145-2C11) from BioLegend. For myeloid cell identification, $CD45^+CD11c^-CD11b^{hi}$ singlets were gated for neutrophils ($Ly6G^+Ly6C^{lo}$), inflammatory monocytes ($Ly6C^{hi}Ly6G^-$), or macrophages ($Ly6C^{lo}Ly6G^-$). For lymphocyte analysis, B cells were identified from the $CD45^+CD19^+$ population, and T cells were identified from the $CD45^+CD3^+$ population that was further gated for $CD4^+$ and $CD8^+$. To determine lymphocyte activation, CD69 antibody was used (H1.2F3, BD). Gating was based on fluorescent-minus-one and isotype control samples. Surface staining was performed in FACS buffer (PBS with 2% BSA) and fixed in 1% paraformaldehyde. Samples were run on a BD LSRII flow cytometer and analyzed through FlowJo Tree Star software.

ELISA

The levels of IL-1β, TNFα, and IL-6 in supernatants from cells or spleens were determined using ELISA kits according to the manufacturer's instructions (R&D Systems). In vitro or in vivo IL-18 level was measured by an ELISA kit from Invitrogen following the manufacturer's instructions.

Statistical analysis

Statistical analyses were performed in Prism 9 (GraphPad) software. In vitro assays were analyzed by unpaired two-tailed Student's *t*-tests for two groups, or ANOVA followed by Bonferroni's post-hoc test for three groups or more. For kinetic cell death assays, the area under the curve (AUC) for each sample was calculated, followed by statistical analyses mentioned above to determine significance. Non-normally distributed data sets were analyzed by Mann-Whitney U tests for comparison between two groups, and Kruskal–Wallis with Dunn's post-hoc tests for three groups or more. Survival curves were compared using log-rank Mantel-Cox tests. Values where $p<0.05$ were considered significant (*$p\leq0.05$, **$p\leq0.01$, ***$p\leq0.001$).

Ethics statement

All animal studies were performed in compliance with the federal regulations set forth in the Animal Welfare Act (AWA), the recommendations in the Guide for the Care and Use of Laboratory Animals of the National Institutes of Health, and the guidelines of the UMass Chan Medical School Institutional Animal Care and Use Committee (IACUC). All protocols used in this study were approved by the UMass Chan Medical School IACUC (protocol A-202000133).

| Table S1. Primers used in quantitative real-time PCR. | | |
|---|---|---|
| **Name** | **F** | **R** |
| *Actin* | CGAGGCCCAGAGCAAGAGAG | CGGTTGGCCTTAGGGTTCAG |
| *Gapdh* | GTCATCATCTCCGCCCCTTCTGC | GATGCCTGCTTCACCACCTTCTTG |
| *Ripk1* Exon4-Alt exon | CCTGAGAATATCCTCGTTGATCGT | GGTTGCTGCTACTTGTTTTGAGTTG |
| *Ripk1* Alt Exon-Exon5 | ACAGCCATCGTCCTCTGGAT | ATGTCTTAAAGGAAGCCACACCAA |
| *Ripk1* Exon9 | TCCTGGTTTTCTTCCTCCCC | CTGTGCAAAGGGGTCATGAG |
| *Clec4e* Exon1-Exon3 | CAAATCGCCTGCATCCCAC | TTCTTGACTGAACCTGATGCCT |
| *Clec4e* Exon1-Exon2 | CAAATCGCCTGCATCCCAC | CGACACATCTGGTGATGAAACAG |
| *Clec4e* Intron2 | GTGTCCCGATGCTGCTGGTA | ACATAGCCGACTTGATCCCA |
| *Clec4e* Exon2-Intron2 | CAGATGTGTCGGTGGGTTCT | GAAAGGAAGCAGGTGGGGTT |
| *Clec4e* Exon3 | CCAGATGTGTCGTAACATATCGC | TTCTTGACTGAACCTGATGCCT |
| *Tnf-α* | CCTATGTCTCAGCCTCTTCT | TTGAGAAGATGATCTGAGTGTG |
| *Il-1β* | TTGACAGTGATGAGAATGACC | CACAGCCACAATGAGTGATA |
| *Raver1* | CTGGCTTCAGTGATGTGGATGC | CATCTCTGCTGTCTCGTACTCC |
| *Ptbp1* | AAGTTTGGCACCGTCCTGAA | CTGGTCTAGTGAAGGCTGGC |

| Table S2. sgRNA sequences used to knock out the indicated genes. | |
|---|---|
| *Raver1_sg1* | CGCCGCAAGATACTGATCCG |
| *Raver1_sg2* | TCCACATCACTGAAGCCAGG |
| *Ptbp1_sg1* | GGTTAACTACTATACATCGG |
| *Ptbp1_sg2* | TCAGTACAGCATCTTACCAA |
| *Ripk1_sg1* | GGCCTCGGGAGCGCGGGTGT |
| *Casp8_sg1* | GATTATGAAAGATCAAGCAC |
| *Clec4e_sg1* | TATCGTCCAGGAGAGCACTT |
| *Clec4e_sg2* | CCTGGTGGTTATCGACACAC |
| *hRAVER1_sg1* | CGCCGCAAGATACTGATCCG |
| *hRAVER1_sg2* | CATCCGTCCAGTGCACGTAG |

### References

1. D. Weng *et al.*, Caspase-8 and RIP kinases regulate bacteria-induced innate immune responses and cell death. *Proc Natl Acad Sci U S A* **111**, 7391-7396 (2014).
2. D. Ratner *et al.*, The Yersinia pestis Effector YopM Inhibits Pyrin Inflammasome Activation. *PLoS Pathog* **12**, e1006035 (2016).
3. G. I. Vladimer *et al.*, The NLRP12 inflammasome recognizes Yersinia pestis. *Immunity* **37**, 96-107 (2012).
4. P. Orning *et al.*, Pathogen blockade of TAK1 triggers caspase-8-dependent cleavage of gasdermin D and cell death. *Science* **362**, 1064-1069 (2018).
5. W. J. Kaiser *et al.*, RIP3 mediates the embryonic lethality of caspase-8-deficient mice. *Nature* **471**, 368-372 (2011).
6. K. Newton, X. Sun, V. M. Dixit, Kinase RIP3 is dispensable for normal NF-kappa Bs, signaling by the B-cell and T-cell receptors, tumor necrosis factor receptor 1, and Toll-like receptors 2 and 4. *Mol Cell Biol* **24**, 1464-1469 (2004).
7. N. Kayagaki *et al.*, Caspase-11 cleaves gasdermin D for non-canonical inflammasome signalling. *Nature* **526**, 666-671 (2015).
8. A. Polykratis *et al.*, Cutting edge: RIPK1 Kinase inactive mice are viable and protected from TNF-induced necroptosis in vivo. *J Immunol* **193**, 1539-1543 (2014).
9. S. B. Berger *et al.*, Cutting Edge: RIP1 kinase activity is dispensable for normal development but is a key regulator of inflammation in SHARPIN-deficient mice. *J Immunol* **192**, 5476-5480 (2014).
10. Y. Wang *et al.*, Chemotherapy drugs induce pyroptosis through caspase-3 cleavage of a gasdermin. *Nature* **547**, 99-103 (2017).
11. J. M. Murphy *et al.*, The pseudokinase MLKL mediates necroptosis via a molecular switch mechanism. *Immunity* **39**, 443-453 (2013).
12. S. Huang *et al.*, Immune response in mice that lack the interferon-gamma receptor. *Science* **259**, 1742-1745 (1993).
13. J. J. Peschon *et al.*, TNF receptor-deficient mice reveal divergent roles for p55 and p75 in several models of inflammation. *J Immunol* **160**, 943-952 (1998).
14. Y. M. Hsu *et al.*, The adaptor protein CARD9 is required for innate immune responses to intracellular pathogens. *Nat Immunol* **8**, 198-205 (2007).
15. R. J. Platt *et al.*, CRISPR-Cas9 knockin mice for genome editing and cancer modeling. *Cell* **159**, 440-455 (2014).
16. R. A. Aglietti *et al.*, GsdmD p30 elicited by caspase-11 during pyroptosis forms pores in membranes. *Proc Natl Acad Sci U S A* **113**, 7858-7863 (2016).
17. S. W. Montminy *et al.*, Virulence factors of Yersinia pestis are overcome by a strong lipopolysaccharide response. *Nat Immunol* **7**, 1066-1073 (2006).
18. S. B. Berger *et al.*, Characterization of GSK'963: a structurally distinct, potent and selective inhibitor of RIP1 kinase. *Cell Death Discov* **1**, 15009 (2015).
19. W. J. Kaiser *et al.*, Toll-like receptor 3-mediated necrosis via TRIF, RIP3, and MLKL. *J Biol Chem* **288**, 31268-31279 (2013).
20. Y. Dondelinger *et al.*, MK2 phosphorylation of RIPK1 regulates TNF-mediated cell death. *Nat Cell Biol* **19**, 1237-1247 (2017).
21. F. C. Peritore-Galve *et al.*, Glucosyltransferase-dependent and independent effects of Clostridioides difficile toxins during infection. *PLoS Pathog* **18**, e1010323 (2022).

22. Y. Chen, X. Lei, Z. Jiang, K. A. Fitzgerald, Cellular nucleic acid-binding protein is essential for type I interferon-mediated immunity to RNA virus infection. *Proc Natl Acad Sci U S A* **118** (2021).
23. L. W. Peterson *et al.*, RIPK1-dependent apoptosis bypasses pathogen blockade of innate signaling to promote immune defense. *J Exp Med* **214**, 3171-3182 (2017).
24. M. Simonet, S. Falkow, Invasin expression in Yersinia pseudotuberculosis. *Infect Immun* **60**, 4414-4417 (1992).
25. Y. Zhang, J. B. Bliska, YopJ-promoted cytotoxicity and systemic colonization are associated with high levels of murine interleukin-18, gamma interferon, and neutrophils in a live vaccine model of Yersinia pseudotuberculosis infection. *Infect Immun* **78**, 2329-2341 (2010).
26. G. Denecker *et al.*, Effect of low- and high-virulence Yersinia enterocolitica strains on the inflammatory response of human umbilical vein endothelial cells. *Infect Immun* **70**, 3510-3520 (2002).
27. D. Ratner *et al.*, Manipulation of Interleukin-1beta and Interleukin-18 Production by Yersinia pestis Effectors YopJ and YopM and Redundant Impact on Virulence. *J Biol Chem* **291**, 9894-9905 (2016).
28. M. Karmakar *et al.*, Neutrophil IL-1beta processing induced by pneumolysin is mediated by the NLRP3/ASC inflammasome and caspase-1 activation and is dependent on K+ efflux. *J Immunol* **194**, 1763-1775 (2015).
29. T. Espevik, J. Nissen-Meyer, A highly sensitive cell line, WEHI 164 clone 13, for measuring cytotoxic factor/tumor necrosis factor from human monocytes. *J Immunol Methods* **95**, 99-105 (1986).
30. P. Mirandola *et al.*, Anticancer agents sensitize osteosarcoma cells to TNF-related apoptosis-inducing ligand downmodulating IAP family proteins. *Int J Oncol* **28**, 127-133 (2006).
31. W. S. Wu *et al.*, Promyelocytic leukemia protein sensitizes tumor necrosis factor alpha-induced apoptosis by inhibiting the NF-kappaB survival pathway. *J Biol Chem* **278**, 12294-12304 (2003).
32. L. Gitlin *et al.*, Essential role of mda-5 in type I IFN responses to polyriboinosinic:polyribocytidylic acid and encephalomyocarditis picornavirus. *Proc Natl Acad Sci U S A* **103**, 8459-8464 (2006).
33. V. A. Rathinam *et al.*, TRIF licenses caspase-11-dependent NLRP3 inflammasome activation by gram-negative bacteria. *Cell* **150**, 606-619 (2012).
34. O. Shalem *et al.*, Genome-scale CRISPR-Cas9 knockout screening in human cells. *Science* **343**, 84-87 (2014).
35. J. G. Doench *et al.*, Rational design of highly active sgRNAs for CRISPR-Cas9-mediated gene inactivation. *Nat Biotechnol* **32**, 1262-1267 (2014).
36. J. G. Doench *et al.*, Optimized sgRNA design to maximize activity and minimize off-target effects of CRISPR-Cas9. *Nat Biotechnol* **34**, 184-191 (2016).
37. B. Wang *et al.*, Integrative analysis of pooled CRISPR genetic screens using MAGeCKFlute. *Nat Protoc* **14**, 756-780 (2019).
38. O. Yukselen, O. Turkyilmaz, A. R. Ozturk, M. Garber, A. Kucukural, DolphinNext: a distributed data processing platform for high throughput genomics. *BMC Genomics* **21**, 310 (2020).
39. A. Dobin *et al.*, STAR: ultrafast universal RNA-seq aligner. *Bioinformatics* **29**, 15-21 (2013).

40. S. Shen *et al.*, rMATS: robust and flexible detection of differential alternative splicing from replicate RNA-Seq data. *Proc Natl Acad Sci U S A* **111**, E5593-5601 (2014).
