## Supplemental figures for "Raver1 links *Ripk1* RNA splicing to caspase-8-mediated pyroptotic cell death, inflammation, and pathogen resistance"

**Figure S1**

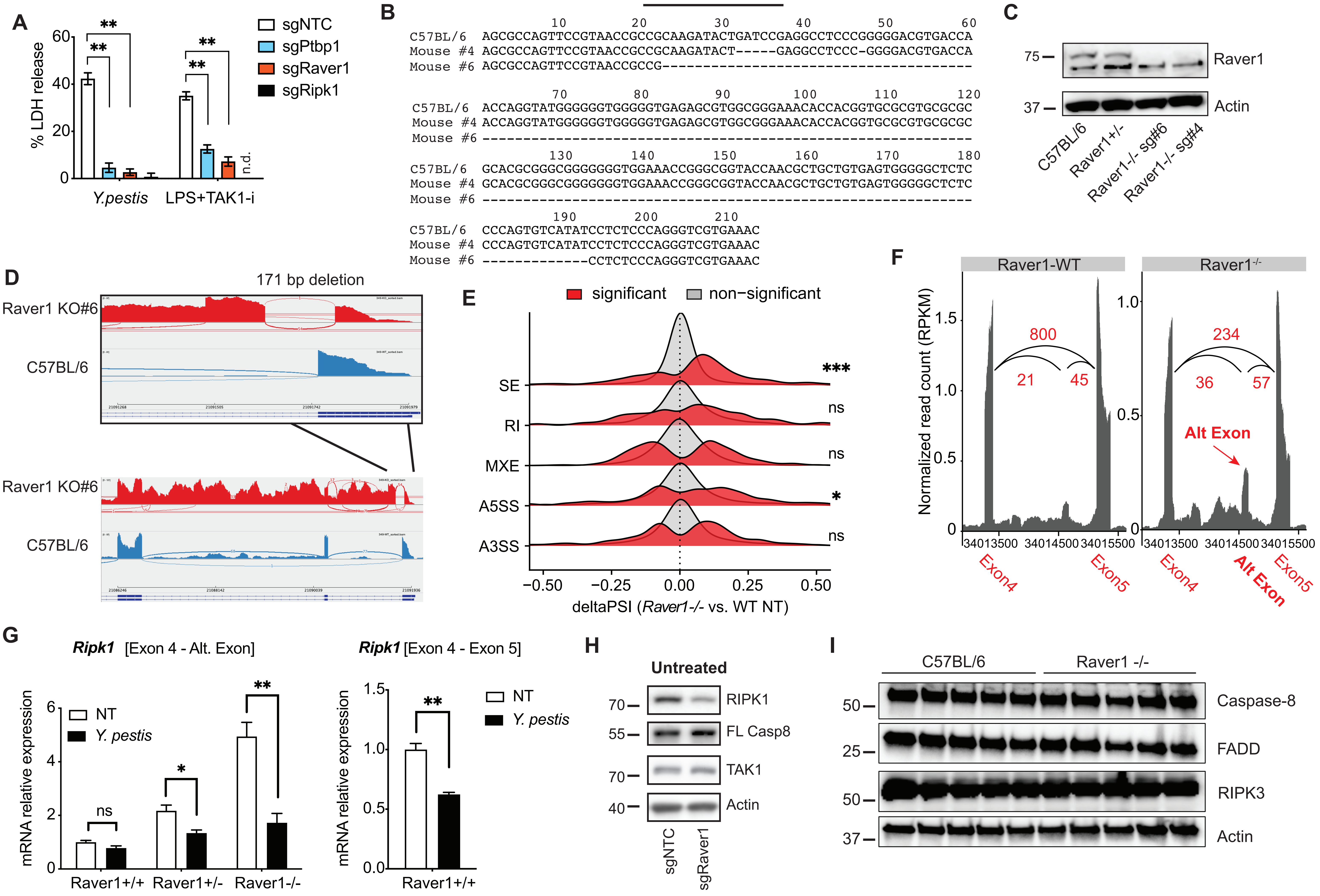
A
% LDH release
60
40
20
0
**
**
**
**
n.d.
Y.pestis
LPS+TAK1-i
sgNTC
sgPtbp1
sgRaver1
sgRipk1
B
sgRNA
10 20 30 40 50 60
C57BL/6 AGCGCCAGTTCCGTAACCGCCGCAAGATACTGATCCGAGGCCTCCCGGGGGACGTGACCA
Mouse #4 AGCGCCAGTTCCGTAACCGCCGCAAGATACT-----GAGGCCTCCC-GGGGACGTGACCA
Mouse #6 AGCGCCAGTTCCGTAACCGCCG--------------------------------------
70 80 90 100 110 120
C57BL/6 ACCAGGTATGGGGGGTGGGGGTGAGAGCGTGGCGGGAAACACCACGGTGCGCGTGCGCGC
Mouse #4 ACCAGGTATGGGGGGTGGGGGTGAGAGCGTGGCGGGAAACACCACGGTGCGCGTGCGCGC
Mouse #6 ------------------------------------------------------------
130 140 150 160 170 180
C57BL/6 GCACGCGGGCGGGGGGGTGGAAACCGGGCGGTACCAACGCTGCTGTGAGTGGGGGCTCTC
Mouse #4 GCACGCGGGCGGGGGGGTGGAAACCGGGCGGTACCAACGCTGCTGTGAGTGGGGGCTCTC
Mouse #6 ------------------------------------------------------------
190 200 210
C57BL/6 CCCAGTGTCATATCCTCTCCCAGGGTCGTGAAAC
Mouse #4 CCCAGTGTCATATCCTCTCCCAGGGTCGTGAAAC
Mouse #6 -------------CCTCTCCCAGGGTCGTGAAAC
C
75
37
Raver1
Actin
C57BL/6
Raver1+/-
Raver1-/- sg#6
Raver1-/- sg#4
D
171 bp deletion
Raver1 KO#6
C57BL/6
Raver1 KO#6
C57BL/6
E
significant
non−significant
SE
RI
MXE
A5SS
A3SS
***
ns
ns
*
ns
−0.50 −0.25 0.00 0.25 0.50
deltaPSI (Raver1-/- vs. WT NT)
F
Raver1-WT
Raver1-/-
Normalized read count (RPKM)
1.5
1.0
0.5
0
800
21
45
234
36
57
Alt Exon
34013500 34014500 34015500
Exon4
Exon5
Alt Exon
G
Ripk1 [Exon 4 - Alt. Exon]
mRNA relative expression
6
4
2
0
NT
Y. pestis
ns
*
**
Raver1+/+
Raver1+/-
Raver1-/-
Ripk1 [Exon 4 - Exon 5]
1.5
1.0
0.5
0.0
**
Raver1+/+
H
Untreated
70
55
70
40
RIPK1
FL Casp8
TAK1
Actin
sgNTC
sgRaver1
I
C57BL/6
Raver1 -/-
50
25
50
37
Caspase-8
FADD
RIPK3
Actin

**Figure S2.**

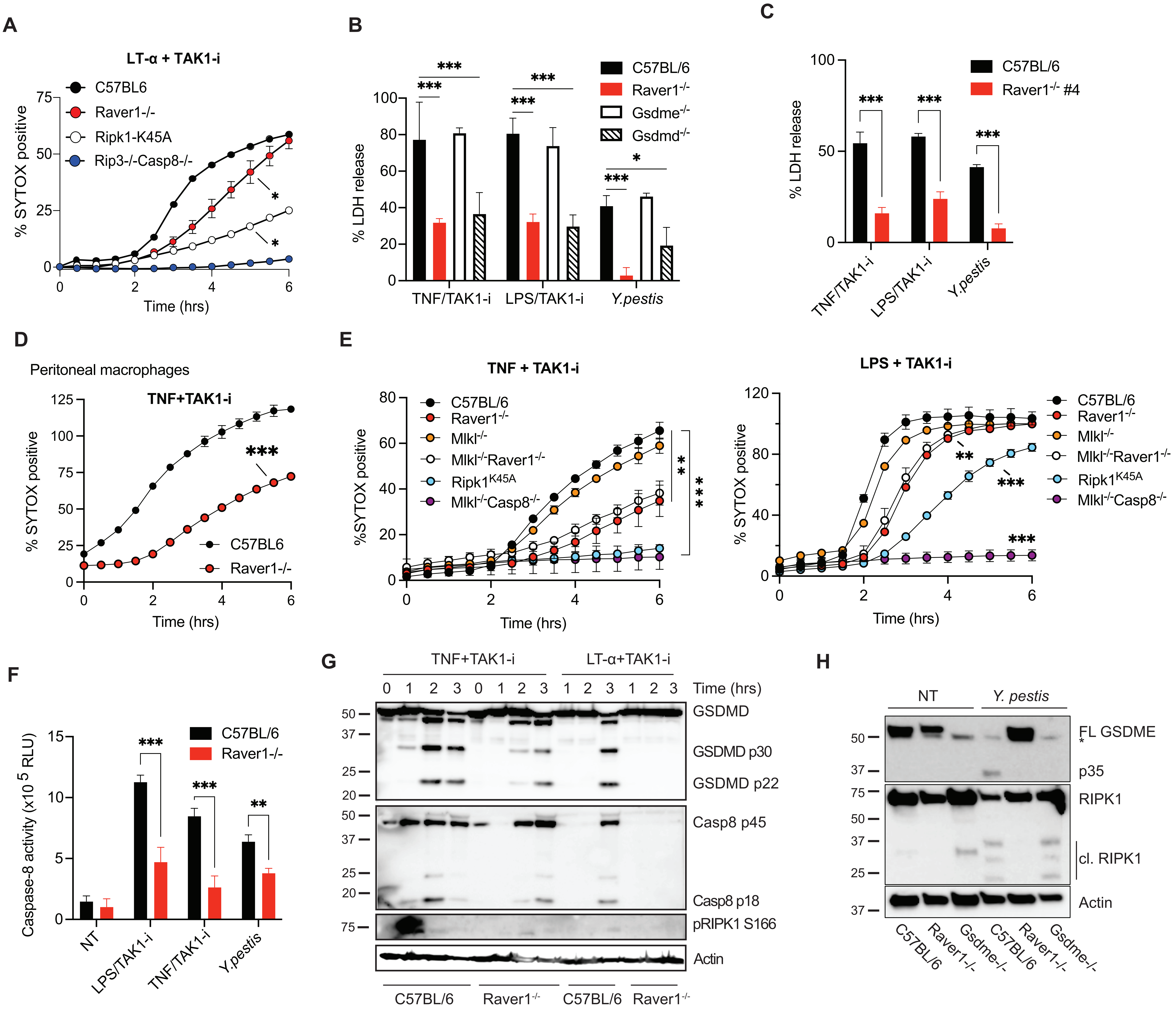
A
LT-α + TAK1-i
C57BL6
Raver1-/-
Ripk1-K45A
Rip3-/-Casp8-/-
% SYTOX positive
Time (hrs)
B
% LDH release
C57BL/6
Raver1-/-
Gsdme-/-
Gsdmd-/-
TNF/TAK1-i
LPS/TAK1-i
Y.pestis
C
% LDH release
C57BL/6
Raver1-/- #4
TNF/TAK1-i
LPS/TAK1-i
Y.pestis
D
Peritoneal macrophages
TNF+TAK1-i
% SYTOX positive
C57BL6
Raver1-/-
Time (hrs)
E
TNF + TAK1-i
%SYTOX positive
C57BL/6
Raver1-/-
Mlkl-/-
Mlkl-/-Raver1-/-
Ripk1K45A
Mlkl-/-Casp8-/-
Time (hrs)
LPS + TAK1-i
% SYTOX positive
Time (hrs)
F
Caspase-8 activity (x10 5 RLU)
C57BL/6
Raver1-/-
NT
LPS/TAK1-i
TNF/TAK1-i
Y.pestis
G
TNF+TAK1-i
LT-α+TAK1-i
Time (hrs)
GSDMD
GSDMD p30
GSDMD p22
Casp8 p45
Casp8 p18
pRIPK1 S166
Actin
C57BL/6
Raver1-/-
C57BL/6
Raver1-/-
H
NT
Y. pestis
FL GSDME
p35
RIPK1
cl. RIPK1
Actin
C57BL/6
Raver1-/-
Gsdme-/-
C57BL/6
Raver1-/-
Gsdme-/-

**Figure S3.**

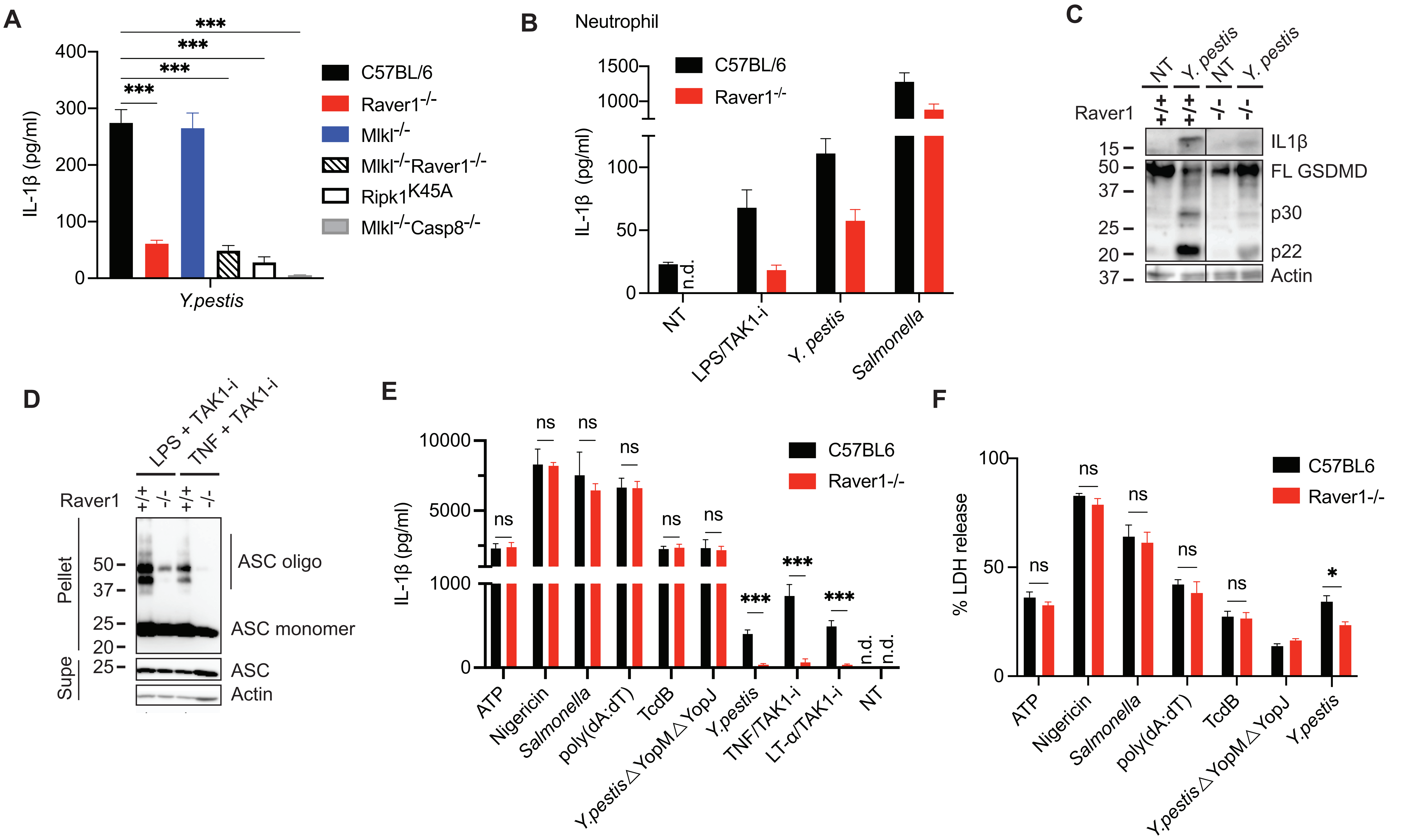

A
***
IL-1β (pg/ml)
Y.pestis
C57BL/6
Raver1-/-
Mlkl-/-
Mlkl-/-Raver1-/-
Ripk1K45A
Mlkl-/-Casp8-/-
B
Neutrophil
IL-1β (pg/ml)
C57BL/6
Raver1-/-
n.d.
NT
LPS/TAK1-i
Y. pestis
Salmonella
C
NT
Y. pestis
Raver1
+/+
-/-
IL1β
FL GSDMD
p30
p22
Actin
D
LPS + TAK1-i
TNF + TAK1-i
Raver1
Pellet
Supe
ASC oligo
ASC monomer
ASC
Actin
E
IL-1β (pg/ml)
C57BL6
Raver1-/-
ns
***
n.d.
ATP
Nigericin
Salmonella
poly(dA:dT)
TcdB
Y.pestis△YopM△YopJ
Y.pestis
TNF/TAK1-i
LT-α/TAK1-i
NT
F
% LDH release
C57BL6
Raver1-/-
ns
*
ATP
Nigericin
Salmonella
poly(dA:dT)
TcdB
Y.pestis△YopM△YopJ
Y.pestis

**Figure S4.**

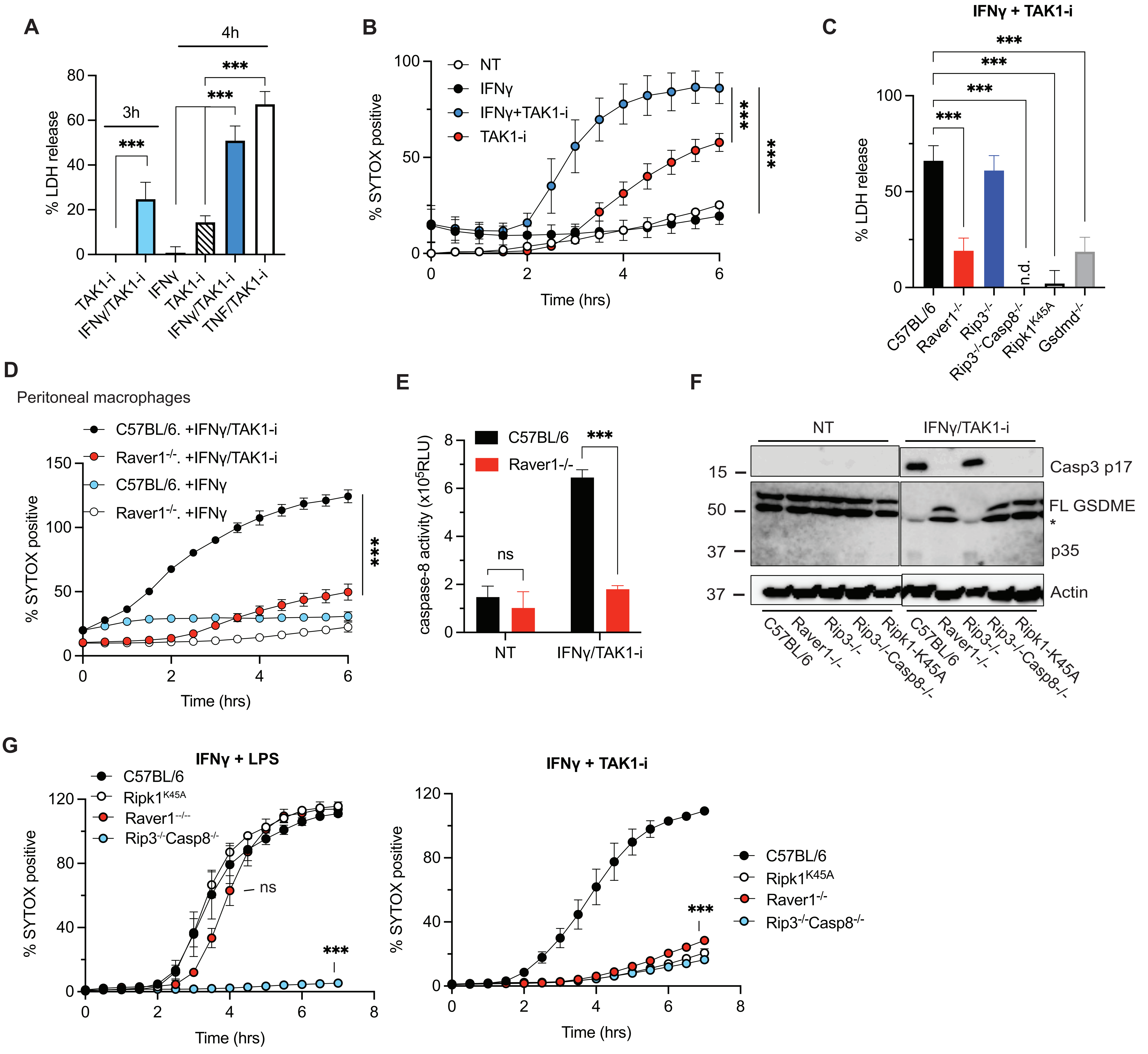

A
3h
4h
% LDH release
TAK1-i
IFNγ/TAK1-i
IFNγ
TAK1-i
IFNγ/TAK1-i
TNF/TAK1-i
B
% SYTOX positive
NT
IFNγ
IFNγ+TAK1-i
TAK1-i
Time (hrs)
C
IFNγ + TAK1-i
% LDH release
n.d.
C57BL/6
Raver1-/-
Rip3-/-
Rip3-/-Casp8-/-
Ripk1K45A
Gsdmd-/-
D
Peritoneal macrophages
C57BL/6. +IFNγ/TAK1-i
Raver1-/-. +IFNγ/TAK1-i
C57BL/6. +IFNγ
Raver1-/-. +IFNγ
% SYTOX positive
Time (hrs)
E
caspase-8 activity (x10^5RLU)
C57BL/6
Raver1-/-
ns
NT
IFNγ/TAK1-i
F
NT
IFNγ/TAK1-i
Casp3 p17
FL GSDME
p35
Actin
C57BL/6
Raver1-/-
Rip3-/-
Rip3-/-Casp8-/-
Ripk1-K45A
G
IFNγ + LPS
C57BL/6
Ripk1K45A
Raver1-/-
Rip3-/-Casp8-/-
% SYTOX positive
ns
Time (hrs)
IFNγ + TAK1-i
Time (hrs)

**Figure S5.**

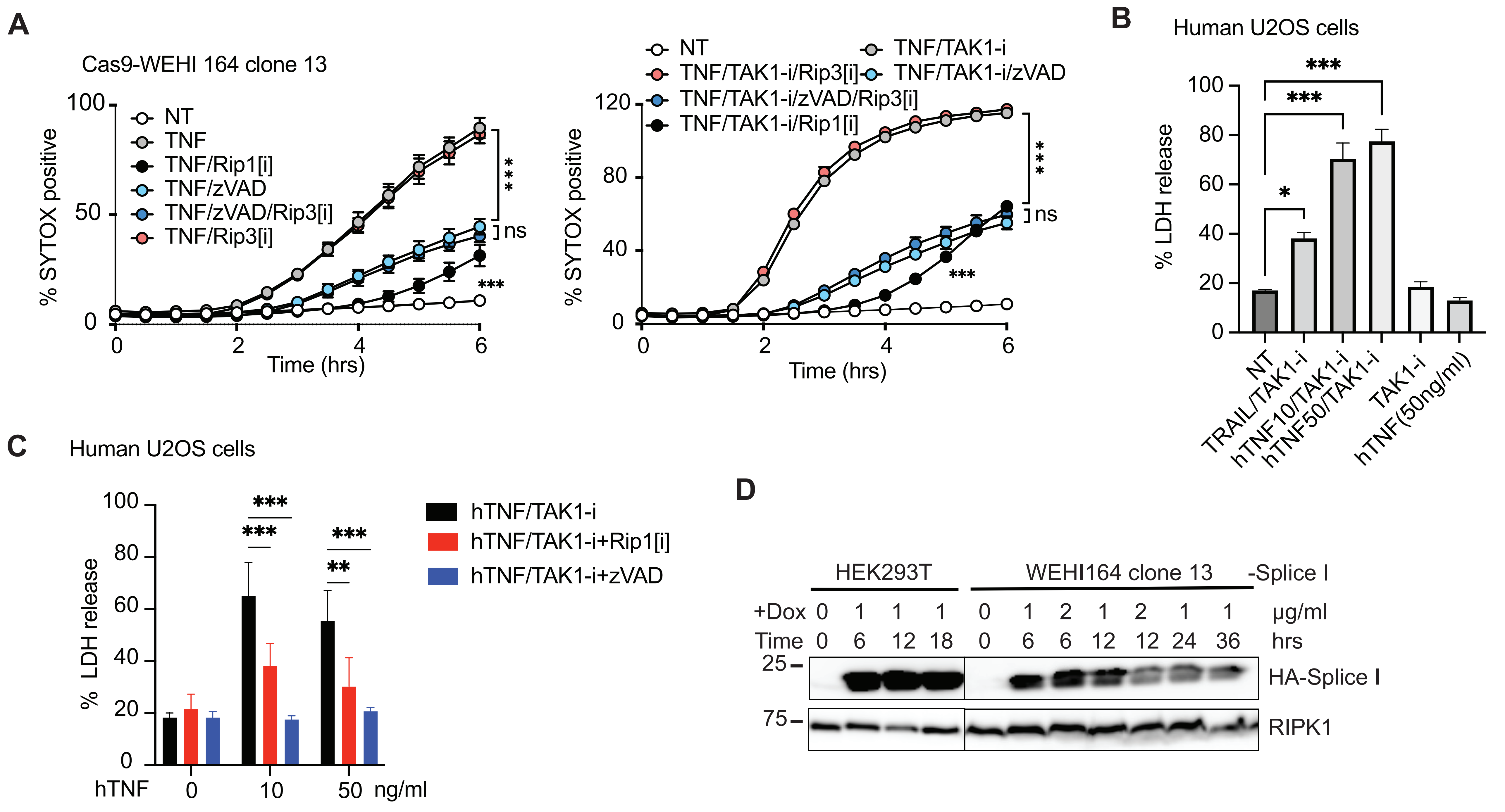
A
Cas9-WEHI 164 clone 13
NT
TNF
TNF/Rip1[i]
TNF/zVAD
TNF/zVAD/Rip3[i]
TNF/Rip3[i]
% SYTOX positive
Time (hrs)
***
ns
***
NT
TNF/TAK1-i
TNF/TAK1-i/Rip3[i]
TNF/TAK1-i/zVAD
TNF/TAK1-i/zVAD/Rip3[i]
TNF/TAK1-i/Rip1[i]
B
Human U2OS cells
% LDH release
NT
TRAIL/TAK1-i
hTNF10/TAK1-i
hTNF50/TAK1-i
TAK1-i
hTNF(50ng/ml)
C
Human U2OS cells
hTNF/TAK1-i
hTNF/TAK1-i+Rip1[i]
hTNF/TAK1-i+zVAD
hTNF 0 10 50 ng/ml
D
HEK293T
WEHI164 clone 13
-Splice I
+Dox 0 1 1 1 0 1 2 1 2 1 1 µg/ml
Time 0 6 12 18 0 6 6 12 12 24 36 hrs
25
75
HA-Splice I
RIPK1

**Figure S6.**

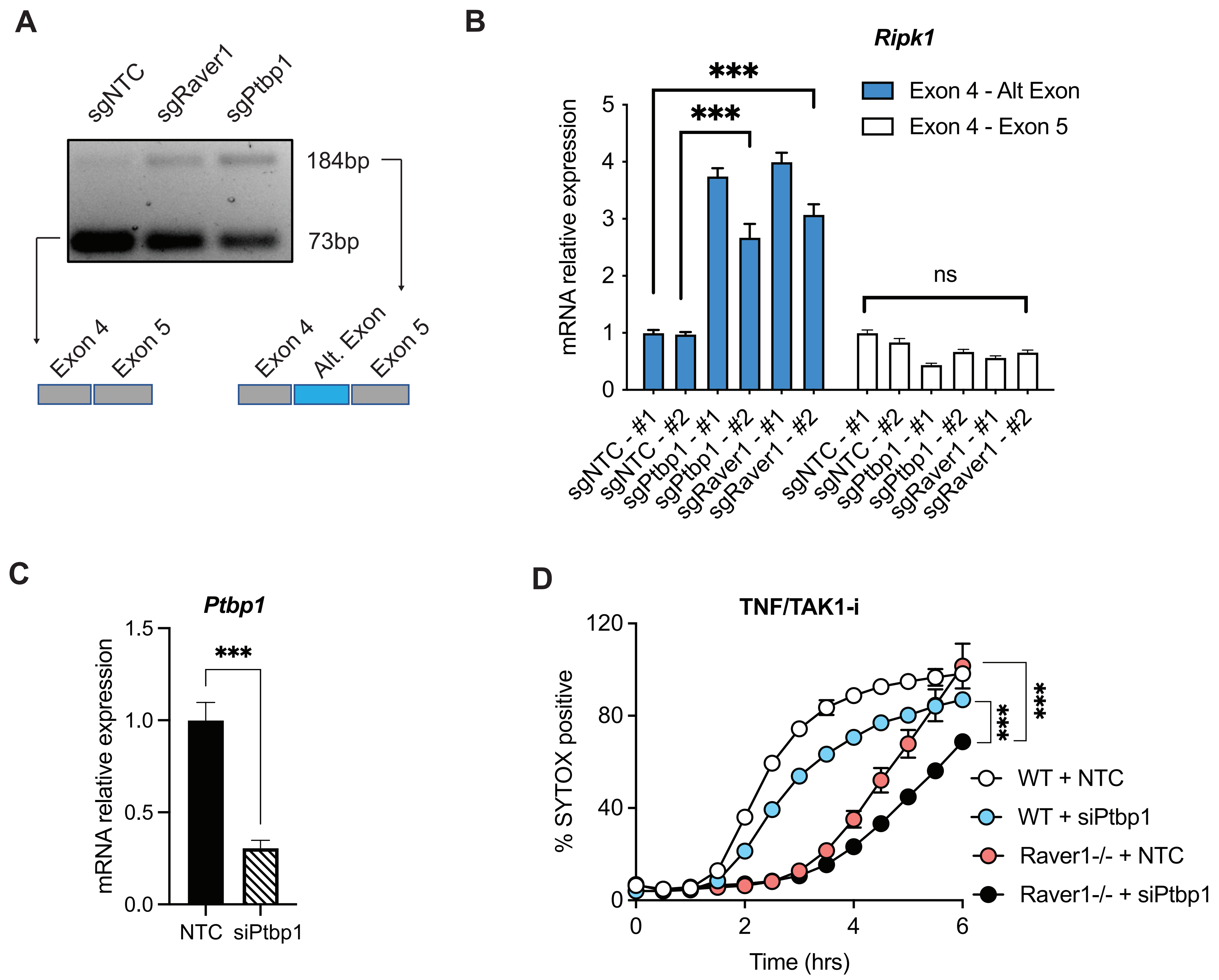
A
sgNTC
sgRaver1
sgPtbp1
184bp
73bp
Exon 4
Exon 5
Exon 4
Alt. Exon
Exon 5
B
Ripk1
Exon 4 - Alt Exon
Exon 4 - Exon 5
***
***
ns
mRNA relative expression
sgNTC - #1
sgNTC - #2
sgPtbp1 - #1
sgPtbp1 - #2
sgRaver1 - #1
sgRaver1 - #2
C
Ptbp1
***
mRNA relative expression
NTC
siPtbp1
D
TNF/TAK1-i
% SYTOX positive
Time (hrs)
***
***
WT + NTC
WT + siPtbp1
Raver1-/- + NTC
Raver1-/- + siPtbp1

**Figure S7.**

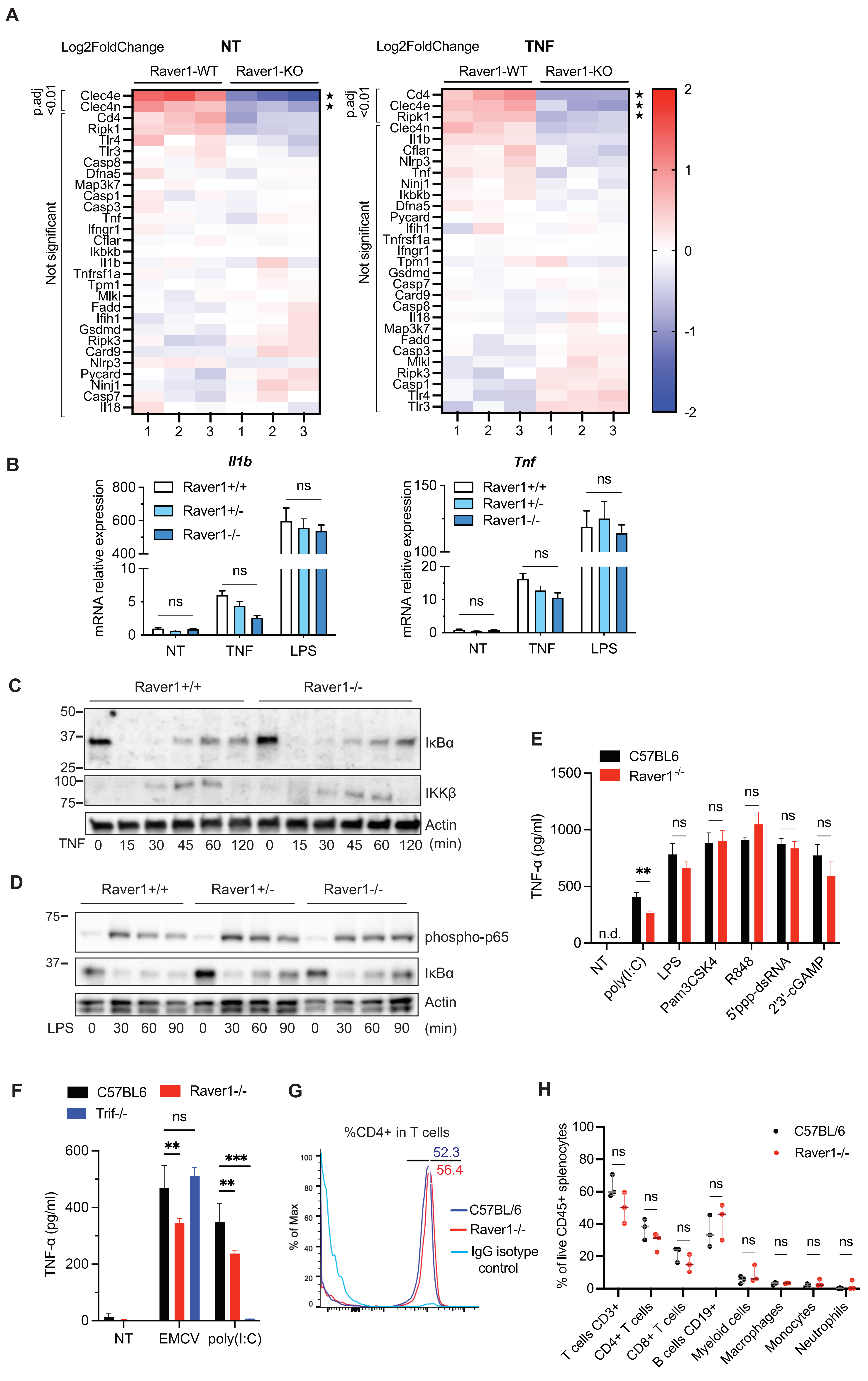
A
Log2FoldChange
NT
Raver1-WT
Raver1-KO
p.adj <0.01
Not significant
Clec4e
Clec4n
Cd4
Ripk1
Tlr4
Tlr3
Casp8
Dfna5
Map3k7
Casp1
Casp3
Tnf
Ifngr1
Cflar
Ikbkb
Il1b
Tnfrsf1a
Tpm1
Mlkl
Fadd
Ifih1
Gsdmd
Ripk3
Card9
Nlrp3
Pycard
Ninj1
Casp7
Il18
TNF
Tpm1
Gsdmd
Card9
Il18
2
1
0
-1
-2
B
Il1b
Tnf
mRNA relative expression
Raver1+/+
Raver1+/-
Raver1-/-
ns
NT
TNF
LPS
C
Raver1+/+
Raver1-/-
IκBα
IKKβ
Actin
TNF 0 15 30 45 60 120 0 15 30 45 60 120 (min)
D
Raver1+/+
Raver1+/-
Raver1-/-
phospho-p65
IκBα
Actin
LPS 0 30 60 90 0 30 60 90 0 30 60 90 (min)
E
C57BL6
Raver1-/-
TNF-α (pg/ml)
n.d.
**
NT
poly(I:C)
LPS
Pam3CSK4
R848
5'ppp-dsRNA
2'3'-cGAMP
F
C57BL6
Raver1-/-
Trif-/-
TNF-α (pg/ml)
**
***
NT
EMCV
poly(I:C)
G
%CD4+ in T cells
52.3
56.4
% of Max
C57BL/6
Raver1-/-
IgG isotype control
H
C57BL/6
Raver1-/-
% of live CD45+ splenocytes
T cells CD3+
CD4+ T cells
CD8+ T cells
B cells CD19+
Myeloid cells
Macrophages
Monocytes
Neutrophils

**Figure S8.**

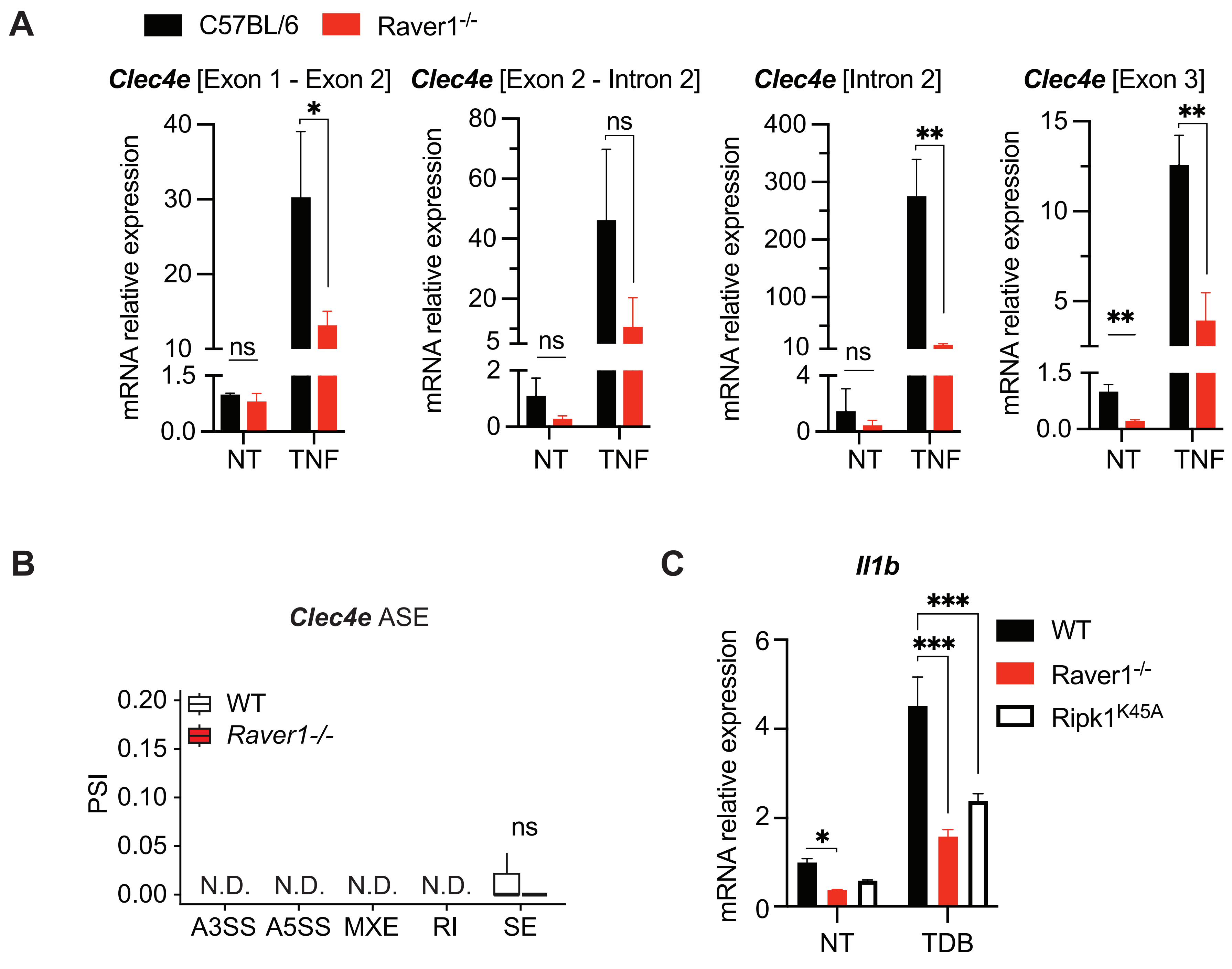
A
C57BL/6
Raver1-/-
Clec4e [Exon 1 - Exon 2]
Clec4e [Exon 2 - Intron 2]
Clec4e [Intron 2]
Clec4e [Exon 3]
mRNA relative expression
ns
*
**
NT
TNF
B
Clec4e ASE
WT
Raver1-/-
PSI
N.D.
A3SS
A5SS
MXE
RI
SE
C
Il1b
WT
Raver1-/-
Ripk1K45A
***
NT
TDB

**Figure S9.**

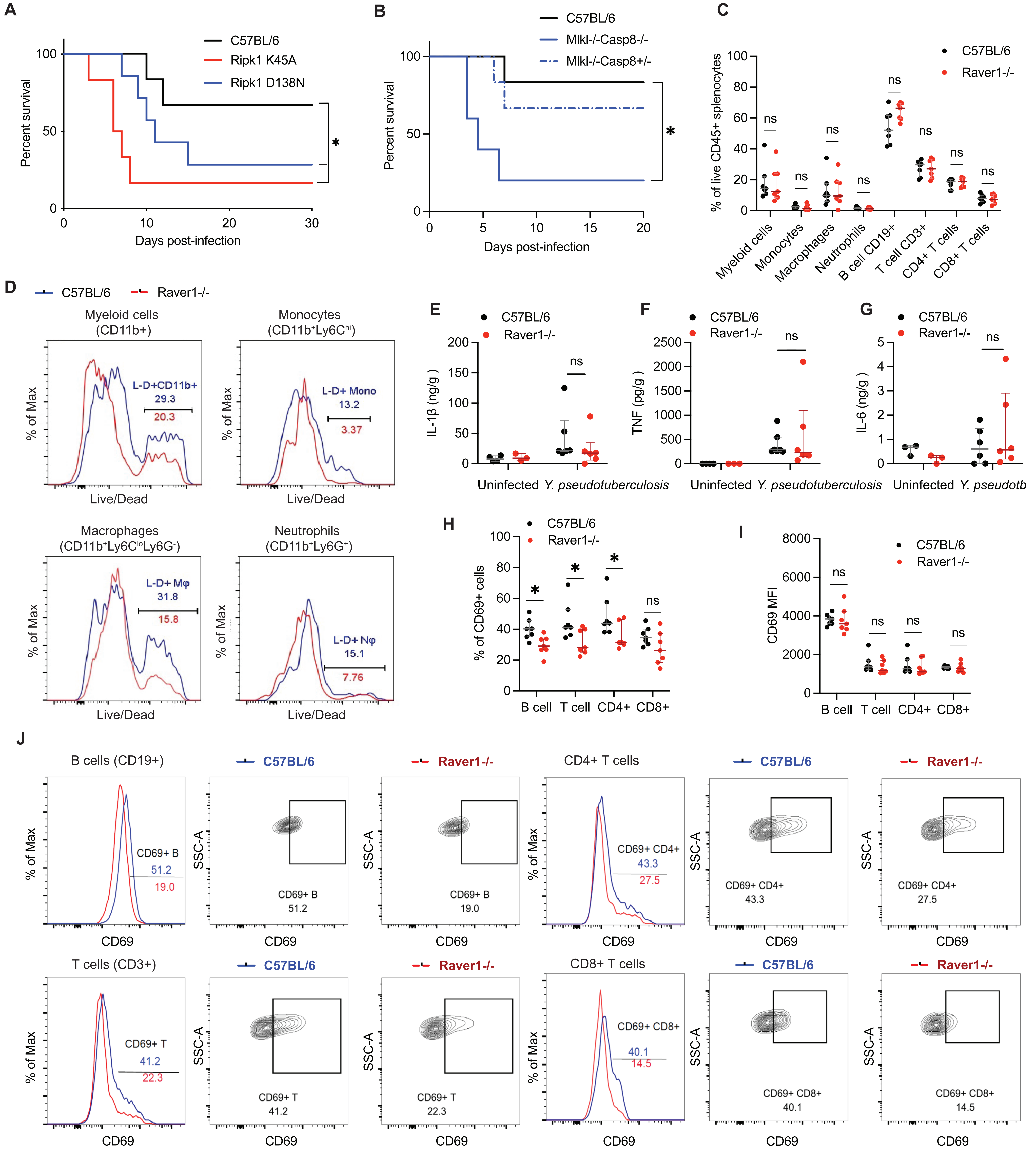
A
C57BL/6
Ripk1 K45A
Ripk1 D138N
Percent survival
Days post-infection
*
B
C57BL/6
Mlkl-/-Casp8-/-
Mlkl-/-Casp8+/-
Percent survival
Days post-infection
*
C
C57BL/6
Raver1-/-
% of live CD45+ splenocytes
Myeloid cells
Monocytes
Macrophages
Neutrophils
B cell CD19+
T cell CD3+
CD4+ T cells
CD8+ T cells
ns
D
C57BL/6
Raver1-/-
Myeloid cells (CD11b+)
L-D+CD11b+
29.3
20.3
Monocytes (CD11b+Ly6Chi)
L-D+ Mono
13.2
3.37
Macrophages (CD11b+Ly6CloLy6G-)
L-D+ Mφ
31.8
15.8
Neutrophils (CD11b+Ly6G+)
L-D+ Nφ
15.1
7.76
% of Max
Live/Dead
E
IL-1β (ng/g)
F
TNF (pg/g)
G
IL-6 (ng/g)
Uninfected
Y. pseudotuberculosis
Y. pseudotb
ns
H
% of CD69+ cells
B cell
T cell
CD4+
CD8+
*
ns
I
CD69 MFI
J
B cells (CD19+)
CD69+ B
51.2
19.0
T cells (CD3+)
CD69+ T
41.2
22.3
CD4+ T cells
CD69+ CD4+
43.3
27.5
CD8+ T cells
CD69+ CD8+
40.1
14.5
C57BL/6
Raver1-/-
% of Max
SSC-A
CD69

**Figure S10.**

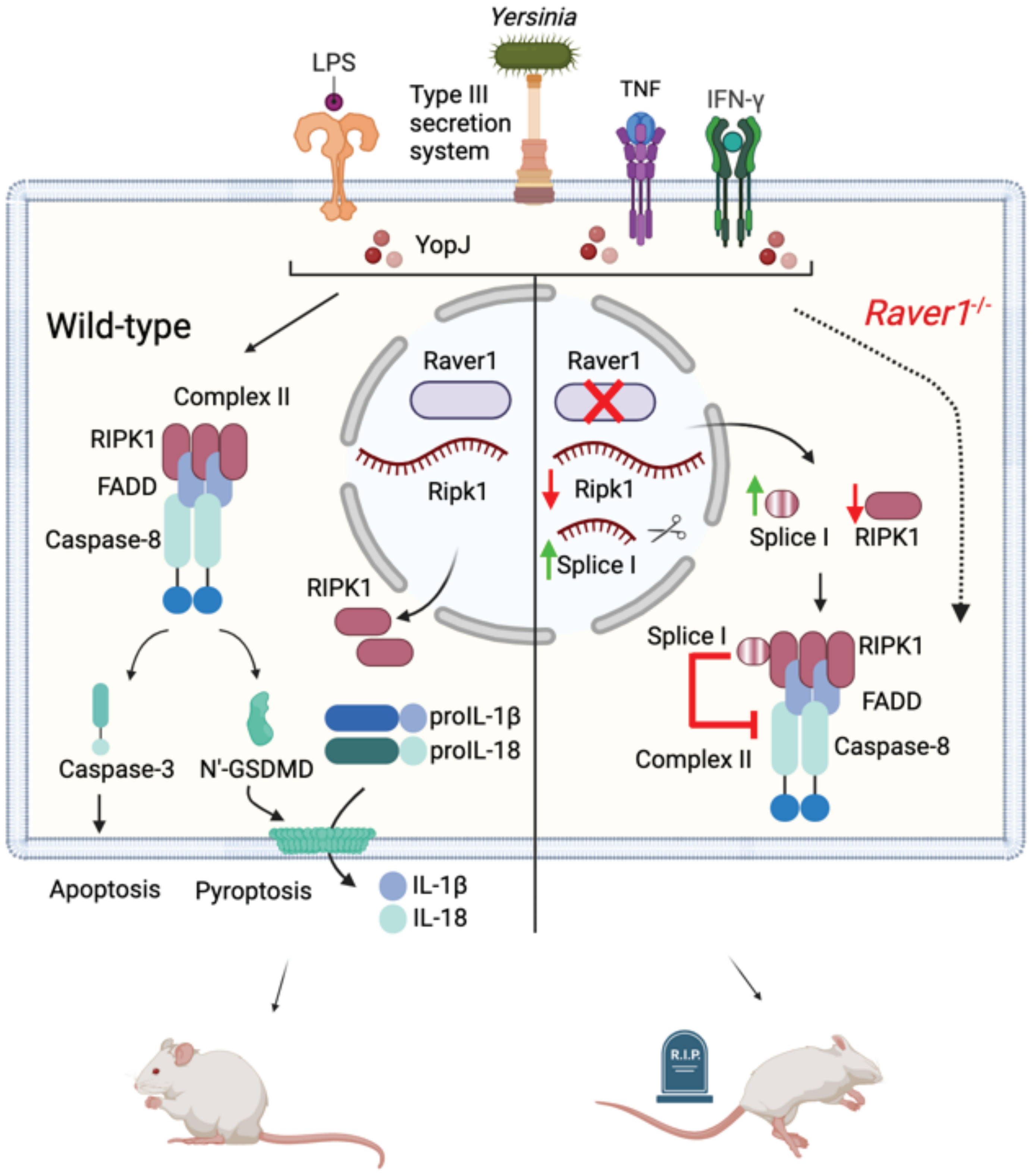


-Resistant to bacterial infection.

-Proper inflammatory responses to eliminate bacteria

-Susceptible to bacterial infection.

-Impaired anti-microbial responses
